# Stochastic spreading models reproduce embolism propagation dynamics in angiosperm xylem networks of vessels connected by bordered pits

**DOI:** 10.1101/2025.10.27.684809

**Authors:** Onerva Korhonen, Steven Jansen, Luciano de Melo Silva, Petri Kiuru, Magdalena Held, Anna Lintunen, Annamari Laurén

**Affiliations:** Tampere University; University of Ulm; University of Graz; University of Eastern Finland; University of Helsinki; University of Eastern Finland / University of Helsinki

**Keywords:** compartment models, SI model, epidemic spreading, directed percolation, plant physiology, xylem hydraulics, xylem networks, embolism spreading, *Betula pendula* Roth, intervessel pit membrane

## Abstract

Plant xylem consists of a network of interconnected vessels, through which water is transported under negative pressure. Filling of vessels with air, or embolism, disturbs this transport process and, in extreme cases, leads to tree mortality. Despite this significance, embolism propagation dynamics are still poorly understood, primarily because xylem is opaque to direct observation. Furthermore, existing models of embolism spreading build excessively on physiological and anatomical parameters, and many misrepresent the inter-vessel pit membrane as a 2D surface. Here, we first extend these physiological models by implementing the pit membrane as a 3D object. Then, we introduce a susceptible-infected (SI) model, a simple stochastic model for tracking spreading through a population, for embolism propagation. After correctly fitting the spreading probability, our SI model reproduces vulnerability curves produced by both the physiological model and empirical data, highlighting that the SI model can address embolism spreading dynamics in plant species, for which detailed physiological data are not available. Furthermore, relating the SI model to the physiological one allows interpreting embolism spreading as a directed percolation process. Elucidating the exact mapping between directed percolation and embolism spreading will likely yield new fundamental insights into the relationships between xylem network architecture and embolism dynamics.

## 1. Introduction

Water is transported in xylem tissue of plants as a continuous column through interconnected dead cells, known as vessels, under negative pressure. This transport is driven by the root-leaf pressure gradient caused by water evaporation at the leaves (Dixon and Joly 1895). However, environmental conditions, in particular prolonged drought, can disturb this transport process by provoking embolism, or filling of xylem vessels with air. As embolism reduces the number of functional xylem vessels, sap transport capacity declines unless new vessels are developed (Tyree and Sperry 1989). Hydraulic failure in xylem, together with carbon starvation, is recognized as the major cause for tree mortality during drought (Adams et al 2017; Anderegg et al 2016; Choat, Brodribb et al 2018; González-Muñoz et al 2018; Mantova et al 2022; McDowell et al 2008; Sevanto et al 2014; Tavares et al 2023).

While the origin of vessel embolism is not fully known, one prominent mechanism is expansion of gas nanobubbles (Ingram et al 2024) that appear in the sap more commonly than traditionally thought (Schenk et al 2015). Embolism events are not randomly distributed among xylem vessels. Instead, embolism is known to spread from embolised vessels to their sap-filled neighbours (Avila, Guan et al 2022; Choat, Badel et al 2016; Guan et al 2021; Kaack, Weber et al 2021; Silva, Pereira et al 2024). However, the field of plant physiology still lacks a golden standard model for embolism spreading that would connect the microscopic function of pit membranes with the mesoscopic xylem structure.

One of the factors that affect embolism spreading in xylem is the interconnectivity between vessels (Loepfe et al 2007). Xylem network models (e.g. Loepfe et al (2007), Mrad, Domec et al (2018), and Mrad, DM Johnson et al (2021)) address this interconnectivity by modelling the xylem as a complex network, or a system of nodes connected by links. Nodes of xylem network models represent the vessels, and links depict the bordered pits between neighbouring vessels. These models build typically on meticulous information about xylem anatomy and physiology, including pit dimensions. For example, Mrad, Domec et al (2018) and Mrad, DM Johnson et al (2021) introduced a model for constructing xylem networks with properties (e.g. average vessel length, vessel density) matching those observed empirically. While this model constructs the xylem network stochastically, the spreading of embolism between neighbouring vessels is deterministic and depends on the bubble propagation pressure (BPP) dictated by intervessel pit membrane properties. Mrad, Domec et al (2018) and Mrad, DM Johnson et al (2021) draw the BPPs from Weibull distribution defined by the largest 10% of measured pore diameters. However, this model does not account for the three-dimensionality of intervessel pit membranes that has important effects on embolism propagation (Kaack, Weber et al 2021; Pereira et al 2023; Zhang et al 2024). As the intervessel pit membrane pores are rather 3D tunnels through the membrane than 2D openings, the resistance of the pit membrane depends on the diameter of the smallest pore constriction instead of the diameter of the pore opening measured at the surface of the pit membrane. However, the 3D aspects of intervessel pit membranes as mesoporous media have not yet been implemented in vessel-level models. Here, we extend the Mrad model by implementing intervessel pits as 3D pit structures (Kaack, Weber et al 2021), where each pore pathway comprises several pore constrictions that together form a channel through the pit membrane.

Physiological models of embolism spreading (e.g. Kaack, Weber et al (2021), Loepfe et al (2007), Lu et al (2025), Mrad, Domec et al (2018), Mrad, DM Johnson et al (2021), and Wason et al (2021)) have made progress in explaining the microscale processes behind embolism spreading. However, the detailed anatomical and physiological data required for constructing these models are available only to a limited number of species, given the arduous measurements required for collecting those data. Further, the physiological models are sensitive to inaccurate assumptions about the complex multiphase interactions between gas, xylem sap with polar lipids, and intervessel pit membranes. To address this problem, we model embolism spreading as a stochastic process, taking place within a complex xylem network, in terms of the susceptible-infected (SI) spreading model. The SI model is a member of the compartment models family (Hethcote 2000; Pastor-Satorras et al 2015). In network science, compartment models have been applied successfully to investigate diverse spreading phenomena, including infectious diseases (Barrat et al 2021; Newman 2002; Rizi et al 2022), rumour spreading (Nekovee et al 2007; Raponi et al 2022; Shah and Zaman 2011), and innovation adoption (Duanmu and Chai 2025). In plant physiology, Roth-Nebelsick (2019) and Torres-Ruiz et al (2016) have introduced compartment models for embolism spreading in a continuous medium. Compared to the physiological embolism spreading models, the main advantage of the SI model is its lightness: instead of the dozen physiological and anatomical parameters that characterise the physiological embolism spreading models, the SI model operates with a single parameter, namely spreading probability (β).

Here, we investigate, to what extent the SI model can replicate embolism spreading produced by a physiological model or extracted from empirical data in the xylem network of *Betula pendula* Roth. We expect that after selecting an optimal spreading probability value for the SI model, the outcomes of the model are comparable to those produced by the physiological model or extracted from empirical data. We assume that this similarity applies both to the percentage loss of conductance (PLC) at the end of the spreading process and to the evolution of hydraulic and network-theoretical properties of the xylem network (e.g. the connected component size and fraction of embolised vessels) during the process. Further, we hypothesize that the vessel pathways in xylem break down faster than the total volume of embolised vessels increases. Thus, we expect that the hydraulic failure related to embolism spreading is primarily caused by xylem network breakdown, rather than the loss of sap in wood tissue. According to this hypothesis, embolism spreading can be interpreted as an instance of a universal network phenomenon, namely directed percolation.

## 2. Materials and methods

### 2.1 Xylem network model

We modelled the xylem anatomy with the network model introduced by Loepfe et al (2007), Mrad, Domec et al (2018), and Mrad, DM Johnson et al (2021). This model represents the xylem as a 3D cylindrical grid of size *N*_*z*_ × *N*_*r*_ × *N*ϕ with the periodic boundary condition 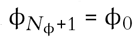, i.e. the number of independent angles considered was *N*ϕ (Fig. 1A-B). The row (*z*), column (*r*), and angular (ϕ) axes of the grid correspond to the axial, radial, and tangential directions of wood tissue.

**Figure 1.**
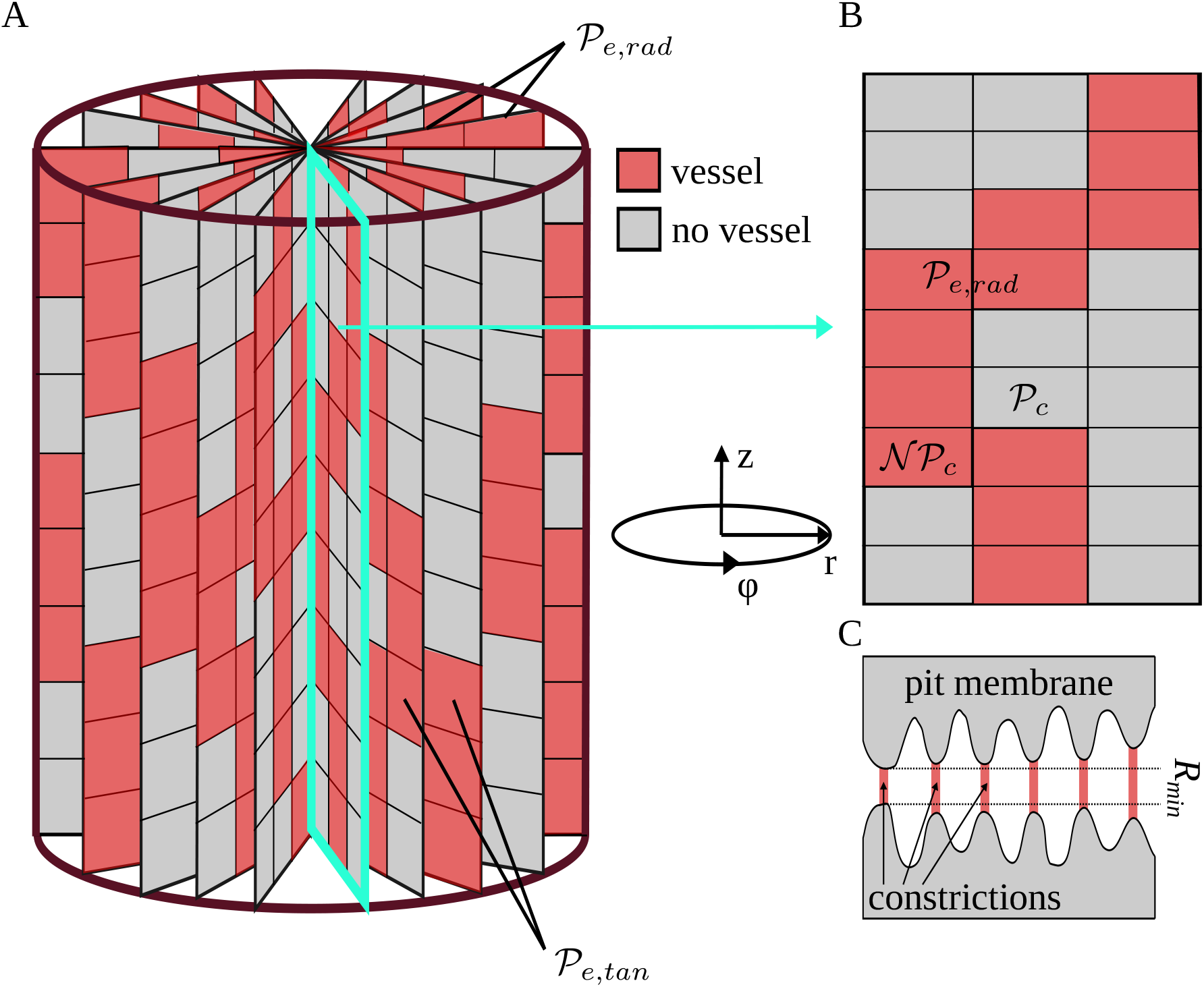
Schematic presentation of the xylem network and pore constriction models. A) The xylem is represented by a 3D cylindrical grid where the row (*r*), column (*z*), and angular (ϕ) dimensions correspond to the axial, radial, and tangential dimensions of a tree. Each grid cell can either belong to a vessel (red cells) or not (grey cells). Neighbouring vessels form intervessel connections in radial direction with probability 𝒫_*e,rad*_ and in tangential direction with probability 𝒫_*e,tan*_. B) In each angular slice of the xylem, each cell above a non-vessel cell starts a new vessel with probability 𝒩𝒫_*c*_, while each cell above a vessel cell ends the existing vessel with probability 𝒫_*c*_. Note that isolated vessels and dead-ends are removed before further analysis. C) Each pit membrane pore is modelled as a set of constrictions (red), forming a channel through the 3D pit membrane (grey). The smallest constriction diameter *R*_*min*_ defines the effective diameter of each pore, and BPP of a pit membrane depends on the largest *R*_*min*_ among all pores of the membrane.

To create the xylem network, we considered the grid cells one by one in the order of their ϕ, *r*, and *z* coordinates. For each cell (*z, r*, ϕ), we checked if the cell could continue an existing vessel, that is, if the corresponding cell at the previous row, (*z* – 1, *r*, ϕ) belonged to a vessel. In this case, the cell (*z, r*, ϕ) ended the vessel (i.e. did not continue it) with probability 𝒫_*c*_ (for all abbreviations and symbols used in the present article, see Table 1). On the other hand, if (*z* – 1, *r*, ϕ) did not belong to an existing vessel, the cell (*z, r*, ϕ) started a new vessel with probability 𝒩𝒫_*c*_. There were two special cases: cells of the first row (i.e. *z* = 0) could not end a vessel and, similarly, cells of the last row (i.e. *z* = *N*_*z*_) could not start a new vessel. Further, all vessels that reached the last row ended automatically independent of the 𝒫_*c*_ parameter. To define vessel diameters, we draw a set of values, corresponding to the number of vessels, from a lognormal distribution, the mean of which was equal to the empirically observed average vessel diameter of *B. pendula* (see Table 2). We assigned the diameters to vessels in relation to vessel length, i.e., so that the longest vessel had the largest diameter. A Python implementation of our network construction model, together with the embolism spreading models and all analysis code used in the present article, is available at https://github.com/onerva-korhonen/tree-percolation.

**Table 1.**
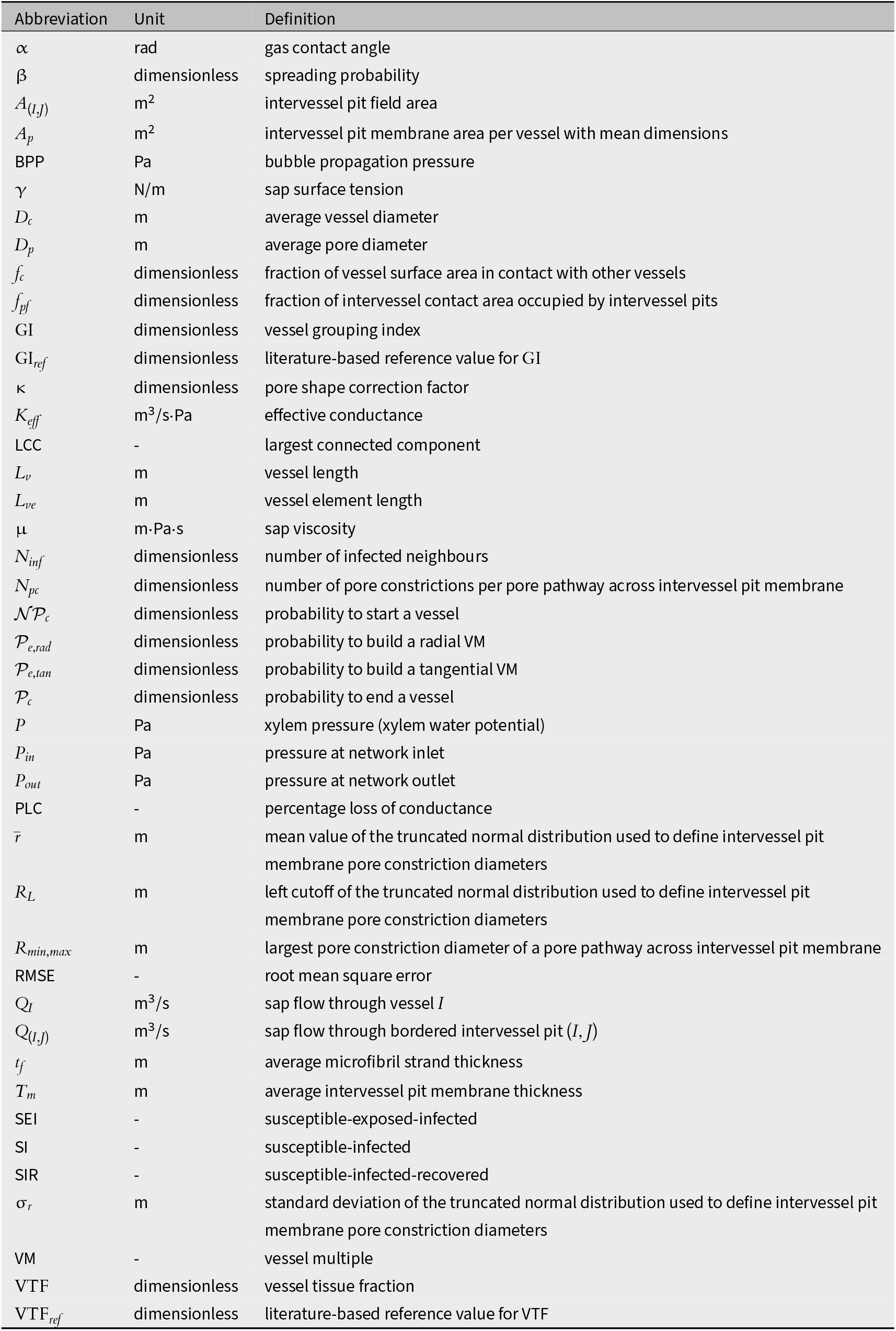
Overview of the abbreviations and symbols used. For values of parameters, see Table 2.

| Abbreviation | Unit | Definition |
| --- | --- | --- |
| $\alpha$ | rad | gas contact angle |
| $\beta$ | dimensionless | spreading probability |
| $A_{(I,J)}$ | m<sup>2</sup> | intervessel pit field area |
| $A_p$ | m<sup>2</sup> | intervessel pit membrane area per vessel with mean dimensions |
| BPP | Pa | bubble propagation pressure |
| $\gamma$ | N/m | sap surface tension |
| $D_c$ | m | average vessel diameter |
| $D_p$ | m | average pore diameter |
| $f_c$ | dimensionless | fraction of vessel surface area in contact with other vessels |
| $f_{pf}$ | dimensionless | fraction of intervessel contact area occupied by intervessel pits |
| GI | dimensionless | vessel grouping index |
| $GI_{ref}$ | dimensionless | literature-based reference value for GI |
| $\kappa$ | dimensionless | pore shape correction factor |
| $K_{eff}$ | m<sup>3</sup>/s·Pa | effective conductance |
| LCC | - | largest connected component |
| $L_v$ | m | vessel length |
| $L_{ve}$ | m | vessel element length |
| $\mu$ | m·Pa·s | sap viscosity |
| $N_{inf}$ | dimensionless | number of infected neighbours |
| $N_{pc}$ | dimensionless | number of pore constrictions per pore pathway across intervessel pit membrane |
| $\mathcal{NP}_c$ | dimensionless | probability to start a vessel |
| $\mathcal{P}_{e,rad}$ | dimensionless | probability to build a radial VM |
| $\mathcal{P}_{e,tan}$ | dimensionless | probability to build a tangential VM |
| $\mathcal{P}_c$ | dimensionless | probability to end a vessel |
| $P$ | Pa | xylem pressure (xylem water potential) |
| $P_{in}$ | Pa | pressure at network inlet |
| $P_{out}$ | Pa | pressure at network outlet |
| PLC | - | percentage loss of conductance |
| $\bar{r}$ | m | mean value of the truncated normal distribution used to define intervessel pit membrane pore constriction diameters |
| $R_L$ | m | left cutoff of the truncated normal distribution used to define intervessel pit membrane pore constriction diameters |
| $R_{min,max}$ | m | largest pore constriction diameter of a pore pathway across intervessel pit membrane |
| RMSE | - | root mean square error |
| $Q_I$ | m<sup>3</sup>/s | sap flow through vessel $I$ |
| $Q_{(I,J)}$ | m<sup>3</sup>/s | sap flow through bordered intervessel pit ($I,J$) |
| $t_f$ | m | average microfibril strand thickness |
| $T_m$ | m | average intervessel pit membrane thickness |
| SEI | - | susceptible-exposed-infected |
| SI | - | susceptible-infected |
| SIR | - | susceptible-infected-recovered |
| $\sigma_r$ | m | standard deviation of the truncated normal distribution used to define intervessel pit membrane pore constriction diameters |
| VM | - | vessel multiple |
| VTF | dimensionless | vessel tissue fraction |
| $VTF_{ref}$ | dimensionless | literature-based reference value for VTF |

**Table 2.** Xylem network and embolism spreading model parameters for *B. pendula*.

| Parameter | Value | Reference |
| --- | --- | --- |
| Vessel dimensions and xylem network properties |  |  |
| $D_c$ | 23.52 $\mu\text{m}$ | Held et al (2025, 2026)<sup>a</sup> |
| $L_{ve}$ | 379 $\mu\text{m}$ | Bhat and Kärkkäinen (1981) |
| $f_c$ | 0.26 | Karimi (2014) |
| $f_{pf}$ | 0.52 | Held et al (2025, 2026) |
| $VTF_{ref}$ | 0.27 | Lintunen and Kalliokoski (2010)<sup>b</sup> |
| $GI_{ref}$ | 2.47 | Alber et al (2019) |
| Intervessel pit and pore properties |  |  |
| $t_f$ | 20 nm | Kaack, Weber et al (2021) |
| $T_m$ | 205 nm | Kaack, Weber et al (2021) |
| $D_p$ | 10.0 nm | Kaack, Weber et al (2021) |
| $\bar{r}$ | 10.0 nm | Kaack, Weber et al (2021) |
| $\sigma_r$ | 7.5 nm | Kaack, Weber et al (2021) |
| $R_L$ | 2.5 nm | Kaack, Weber et al (2021) |
| $N_{pc}$ | 7 | Kaack, Weber et al (2021)<sup>c</sup> |
| $\kappa$ | 0.5 | Kaack, Weber et al (2021) and Schenk et al (2015) |
| $\alpha$ | 0 | Kaack, Weber et al (2021) |
| Sap properties |  |  |
| $\mu$ | 1.002 m·Pa·s | Mrad, Domec et al (2018) |
| $\gamma$ | 25 mN/m | Kaack, Weber et al (2021) |

This procedure yielded a set of vessels with known physical locations. These vessels formed the nodes of the xylem network, while the links of the network corresponded to the connections between neighbouring vessels, known as vessel multiples (VMs). To create these links, we identified the neighbouring cells, or cells sharing a face or a vertex in *r* or ϕ directions, of each cell belonging to a vessel. A VM connected two neighbouring cells in different vessels with probability 𝒫_*e,rad*_ in the radial direction and with probability 𝒫_*e,tan*_ in the tangential direction.

Several VMs can exist between a vessel pair. We considered all VMs when calculating the sap flow through the xylem and the related effective conductance of the xylem network (section 2.2). However, for the sake of simplicity, in the SI model for embolism spreading (section 2.3.2), we considered the vessel network unweighted, meaning that all links have an equal weight, independently of the pit area shared by the vessels connected by the link.

The procedure for creating vessels and VMs may lead to isolated vessels. Such vessels are rare or non-existent in real plants (Bosshard and Kučera 1973; Brodersen et al 2011; Burggraaf 1972). Thus, we considered them artifacts of our network construction algorithm and removed them from any further analysis. Similarly, we removed any dead-ends, that is, vessel groups that were not connected to both the inlet and the outlet rows, following the example of earlier xylem network construction algorithms (Brodersen et al 2011; Mrad, Domec et al 2018). Importantly, our model included only one type of conducting cells, vessels.

### 2.2 Hydraulic properties of the xylem network

We measured the functionality of the xylem network in terms of effective conductance *K*_*eff*_ defined as the total sap flow through the network normalized by the pressure difference between the network inlet and outlet. Following Mrad, Domec et al (2018), we calculated the sap flow *Q*_*I*_ through vessel *I* in terms of the Hagen-Poiseuille equation:

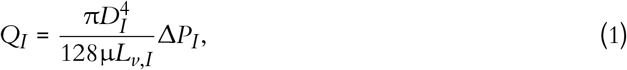

where *D*_*I*_ denotes the diameter and *L*_*v,I*_ the length of vessel *I*, µ is sap viscosity, and Δ*P*_*I*_ is the pressure difference between the ends of the vessel. Further, the flow through VM (*I, J*) is obtained by considering the Sampson flow resistance for pores in a plate and a Poiseuille flow contribution through the finite membrane thickness as:

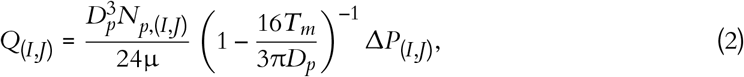

where *D*_*p*_ is the average intervessel pit membrane pore diameter, *T*_*m*_ is the average intervessel pit membrane thickness, and Δ*P*_(*I,J*)_ is the pressure difference over the bordered intervessel pit pair (Dagan et al 1982; Mrad, Domec et al 2018). The number of pores *N*_*p*,(*I,J*)_ is obtained, following Mrad, Domec et al (2018) and Sperry and Hacke (2004), by dividing the VM pit field area *A*_*I,J*_ by the area of an average pit membrane pore:

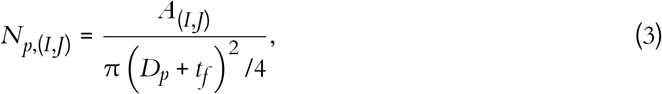

where *t*_*f*_ is microfibril strand thickness. The formula for *A*_(*I,J*)_ is given by Mrad, Domec et al (2018) and Loepfe et al (2007) as

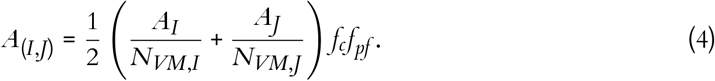

Here, the fraction of the surface area *A*_*I*_ and number of VMs *N*_*VM,I*_ of vessel *I* defines the theoretical vessel surface area available for VM (*I, J*), the vessel contact fraction *f*_*c*_ tells the average fraction of wall area shared with other vessels out of total vessel surface area, and *f*_*pf*_ is the fraction of the contact area between two vessels, *A*_*I*_*f*_*c*_, occupied by intervessel pit membranes.

Combining equations 1 and 2 allowed obtaining the total water flow *Q* through the system, assuming continuity everywhere except for the inlet and outlet and given the inlet and outlet pressures *P*_*in*_ and *P*_*out*_. Finally, from *Q* we obtained the effective conductance as

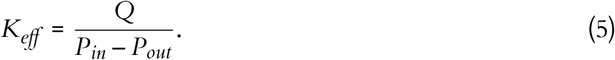

As stated above, the total flow *Q*, and consequently *K*_*eff*_ calculated through equation 5, originate from a combination of Pouseuille and Sampson flows. Both these flows are solutions to the general Navier-Stokes equations for the motion of viscous fluids derived under special conditions. Therefore, in our Python implementation, we simulated the sap flow using the `StokesFlow` algorithm of the OpenPNM Python package (Gostick et al 2016), version 3.4.1.

### 2.3 Embolism spreading

To investigate the vulnerability of the xylem network to embolism, we simulated embolism spreading between vessels with two approaches. First, we considered a *physiological spreading model* based on the pit-level properties of VMs. Second, we showed that the *SI model* (Hethcote 2000; Pastor-Satorras et al 2015) can replicate the results of the physiological spreading model while significantly reducing the amount of free model parameters and thus reducing the model complexity. In both models, as soon as a vessel became embolised, it was removed from the network together with all its VMs, so that it no longer contributed to sap flow or any network properties (see section 2.4). In other words, at any given moment, the intervessel network contained only sap-filled vessels, and thus the number of nodes in the network decreased during the embolism spreading process.

Both models assumed that there is at least one embolised vessel, the spreading seed, in the xylem at the beginning of the embolism spreading process. This assumption is reasonable for xylem tissue of perennial, woody plants because embolism is often observed in older growth rings or primary xylem (Choat, Badel et al 2016). Our models described how this initial embolism spreads in the xylem but did not aim to explain mechanistically how the seed vessel became embolised. Further, we did not consider refilling of vessels as the time scale of refilling is notably slower than the embolism spreading time scales we considered here. Further, while seasonal refilling takes place in *B. pendula* under positive pressures (Strati et al 2003), this recovery mechanism is less probable under normal, negative xylem pressures (Choat, Brodribb et al 2018).

#### 2.3.1 Physiological spreading model

In the physiological spreading model, the spreading of embolism between two vessels depended on the VM-specific BPP. We calculated the BPPs using the pore constriction model introduced by Kaack, Weber et al (2021) (referred to as Model 1 in the original article) (Fig. 1C). In other words, we modelled the intervessel pit membrane as a 3D object so that each pit membrane pore formed a channel, consisting of several constrictions, through the 3D pit membrane. We drew the constriction diameters from a left-truncated normal distribution with mean value 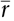, standard deviation σ_*r*_, and cutoff *R*_*L*_, and the smallest constriction diameter defined the diameter of the pore. The number of constrictions per pore depended on the intervessel pit membrane thickness as

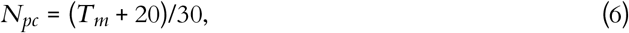

where *T*_*m*_ was expressed in nanometers and a 10 nm spacing between microfibril strands was assumed (Kaack, Weber et al 2021). To define the BPP of an intervessel pit field with area *A*_*p*_, we simulated a set of pores so that their total area was as close as possible to *A*_*p*_ without exceeding it. Calculation of the pit BPP was based on the Young-Laplace equation, a general equation that describes how the pressure difference across the interface between two fluids depends on the interface curvature and surface tension. Following Kaack, Weber et al (2021) and Schenk et al (2015), we calculated BPP using a modified Young-Laplace equation as

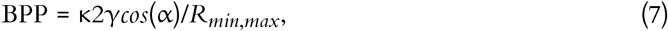

where κ is a dimensionless pore shape correction factor that accounts for irregularities in pore shape, in particular for deviation from the ideal cylindrical shape, for which the modified Young-Laplace equation has originally been derived (Emory 1989; Schenk et al 2015; US EPA Office of Water 2005). Further, α is the gas contact angle between the propagating air bubble and the sap surface and *R*_*min,max*_ the largest pore diameter, that is, the largest among the smallest constriction diameters found in all the pores of the intervessel pit membrane. Finally, γ is the equilibrium value of the dynamic sap surface tension that depends on the local lipid concentration at sap-air interfaces; this value is notably smaller than the surface tension of pure water (Kaack, Altaner et al 2019) (for exact parameter values used, see Table 2). For each pair of neighbouring vessels *I* and *J*, we defined BPP(*I, J*) as the smallest of the BPPs of the VMs connecting *I* and *J*.

When defining the number of pores in a pit membrane, we calculated the area of each pore using the smallest constriction diameter as pore diameter. This gives a theoretical upper limit for the number of pores for a case where all constrictions of each pore have the same, minimum diameter. Similarly, calculating pore area based on the largest constriction diameter gives a theoretical lower limit that corresponds to a case where the largest constrictions of each pore are aligned. In reality, pores have constrictions larger than the smallest one, constrictions of different pores are not aligned, and the pore tunnels may be curved, which makes estimating the actual number of pores per pit challenging. While we assume that the effects of estimating the number of pores in terms of the theoretical upper limit to be small, this decision may have yielded slightly underestimated BPPs and, correspondingly, lower embolism resistance and less negative *P*_50_ values.

Air moves from an embolised vessel to its sap-filled neighbours through a series of snap-off events creating nanobubbles. These bubbles may expand and embolise the sap-filled neighbouring vessel. On the other hand, they may become stabilized, in which case their dynamic surface tension, caused by the stretching of the surfactant molecules coating the bubble, prevents expansion (Kaack, Altaner et al 2019; Schenk et al 2015; Silva, Bujnowski et al 2025). The fracture of this coating typically leads to sudden increase of surface tension and collapse of the nanobubble (Kaack, Altaner et al 2019; Schenk et al 2015; Silva, Bujnowski et al 2025). However, modelling the coating process would require detailed information about the multiphase interactions inside the intervessel pit membrane, including the nanoscale interactions between surfactants, xylem sap, and gas bubbles. For simplicity, we did not model this process explicitly but instead assumed that embolism spreading happens between vessels *I* and *J* every time when the sap pressure exceeds the BPP. Therefore, at each time step *t* of the spreading simulation, we embolised each non-embolised vessel *I* that had an embolised neighbour *J* (i.e. a neighbour *J* with embolisation time < *t*) so that BPP(*I, J*) *≤ P*, where *P* was the sap pressure in the xylem.

#### 2.3.2 SI model

The SI model is a stochastic model that enables tracking spreading through a population straight-forwardly and with a minimum number of free parameters. This model and related compartment models are widely applied to study diverse phenomena including spreading of pathogens (e.g. Barrat et al (2021) and Rizi et al (2022)), propagation of rumours and fake news (e.g. Nekovee et al (2007) and Shah and Zaman (2011)), innovation adoption (e.g. Duanmu and Chai 2025), and spreading of invasive species (e.g. Strickland et al (2015)). The SI model divides the population (here, the xylem vessels) into two compartments: the susceptible (S) compartment, which is vulnerable to an infection but not yet infected, and the infected (I) compartment that contains infected individuals. At each time step, every individual in the S compartment can catch the infection from any of their neighbours in the I compartment with probability β. Since these spreading events are independent, the total infection probability of a susceptible individual with *N*_*inf*_ infected neighbours is

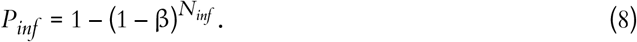

This S-I transition is the only transition between compartments in the SI model. The model does not contain any recovery mechanisms, and thus transitions from the I compartment to the S compartment are not possible.

In our SI model of embolism spreading, the S compartment represented the sap-filled vessels, while the I compartment corresponded to the embolised ones.

#### 2.3.3 Vulnerability curves

To investigate the xylem network’s resistance to embolism under different conditions, we first simulated 100 individual xylem networks starting from a 100 × 10 × 100 grid (i.e. each network has at most 100 × 10 × 100 = 100 000 vessel elements). For each of these networks, we selected the seed vessels, from which the embolism spreading started, by identifying the components (i.e. groups of nodes that were reachable from each other through the network links) of the xylem network and picking a random vessel per component. These seed vessels represent the embolised vessels that are present in real-world xylem (Carmesin et al 2023).

For each of these networks, we simulated embolism spreading with the physiological spreading model for a set of 79 *P* values: 8 values ranging from 0 MPa to -0.875 MPa with the step size of 0.125 MPa, 50 values ranging from -1 MPa to -1.49 MPa with the step size of 0.01 MPa, and final 21 values ranging from -1.5 MPa to -4 MPa with the step size of 0.125 MPa. We selected the uneven step size to emphasize the pressure area around the expected *P*_50_ value (i.e. the pressure where 50% of *K*_*eff*_ is lost). We set the embolisation time of the seed vessels to zero and ran the physiological spreading model for each *P* until the spreading saturated, that is, until the fraction of embolised vessels did not change for 20 time steps. The final *K*_*eff*,*phys*_ for each *P* was averaged across the 100 simulation iterations.

Then, for each *P*, we calculated the PLC by comparing the *K*_*eff*,*phys*_ at spreading saturation point to *K*_*eff*,0_ calculated for the xylem with no embolised vessels. Finally, we constructed the physiological vulnerability curve by drawing the PLC as a function of *P*.

To find the SI model spreading probability β that best approximates embolism spreading at each *P*, we ran for each of the simulated networks the SI model for 65 β values: 30 values ranging from 0 to 0.15 with step size of 0.005 and additional 35 values ranging from 0.175 to 1 with step size of 0.025. We selected the uneven step size through preliminary simulations that used an evenly spaced probability range from 0 to 1 with step size of 0.15. The outcomes of these simulations hinted that lower β values yield a better fit to the physiological spreading model. Thus, we selected for the final analysis an uneven step size that emphasizes the lower β values.

Again, we ran the model until the spreading saturated, and calculated the final *K*_*eff*,*SI*_ as an average across the 100 iterations. Then, we selected for each *P* the β that produced *K*_*eff*,*SI*_ closest to the corresponding *K*_*eff*,*phys*_. Next, we calculated the PLC for the selected β values and drew the SI-based vulnerability curve as a function of *P*.

In addition to optimizing β against the physiological model, we fitted the SI model also against an empirical *B. pendula* vulnerability curve published by González-Muñoz et al (2018). Unlike for the physiological model, the absolute *K*_*eff*_ values were not available for the empirical data. Therefore, we optimized β directly in terms of PLC, selecting for each *P* value, separately, the β value that produced the PLC closest to that extracted from the empirical data.

### 2.4 Network properties

We explored the embolism spreading in xylem further in terms of a set of network properties calculated at each time step of the spreading process. For all these properties, we reported the mean and standard deviation across 100 iterations. Note that as described in Section 2.3, we removed vessels from the intervessel network as soon as they became embolised, and thus only sap-filled vessels contributed to the network properties.

#### Largest connected component (LCC) size

Depending on the values of 𝒫_*c*_, 𝒩𝒫_*c*_, 𝒫_*e,rad*_, and 𝒫_*e,tan*_, the xylem network could be divided into several components. Every vessel of a component was reachable from other vessels of the same component following network links, while there were no links between vessels belonging to different components. The LCC size was defined as the number of vessels in the largest network component. Because of the systematical removal of embolised vessels from the network, the LCC contained sap-filled vessels only.

#### Functional LCC size

We considered a network component functional if it contained at least one vessel starting from the inlet row and at least one vessel ending at the outlet row. In other words, a functional component spanned the whole xylem space in vertical direction and thus contributed to the sap flow through the xylem and *K*_*eff*_. We defined functional LCC size as the number of vessels in the largest functional component. The difference between the LCC and the functional LCC was the sap flow: while both the LCC and the functional LCC consisted of sap-filled vessels, the LCC did not necessarily connect to both network inlet and outlet and thus there was no guaranteed sap flow through it.

*Prevalence* is an epidemiological measure defined as the fraction of infected individuals out of the total population. Here, we calculated prevalence as the number of embolised vessels divided by the total number of vessels in the intact xylem network.

#### Nonfunctional sap volume

Unlike the functional components, nonfunctional xylem network components lacked either the inlet, the outlet, or both. Thus, there was no sap flow through the nonfunctional components. These components arose from embolism spreading: while all components of a xylem network with no embolised vessels were functional and filled with sap, a component could lose its connection to the inlet or the outlet because of embolism. In such a case, the sap in the now nonfunctional component could not exit the xylem because of the lack of either access to outlet or the driving pressure difference between the ends of the component. We defined nonfunctional sap volume as the total volume of this sap trapped in the xylem, which was equal to the total volume of vessels in nonfunctional components.

*Number of inlets and outlets per component* was defined as the average number of vessels starting from the inlet row or ending at the outlet row per component.

The investigated network measures were related to each other. In particular, assuming that embolism spread at the same speed in all network components, we expected the relationship between the LCC size and the functional LCC size to be positive and approximately linear so that the LCC size set the upper boundary for the functional LCC size. We expected both the LCC size and the functional LCC size to decrease during embolism spreading, both due to the removal of embolised vessels from the network and because this removal could divide existing network components into several smaller ones. Further, we expected prevalence to increase approximately linearly with the decreasing LCC size, assuming that embolism events were equally distributed across network components. The expected relationship of the nonfunctional sap volume with the largest LCC size was more complicated. The nonfunctional sap volume increased with the increasing LCC size when more components lost their connection to the inlet or to the outlet. However, the nonfunctional sap volume could also decrease during embolism spreading, as also vessels that belong to nonfunctional components could become embolised. Finally, we expected the number of inlets and outlets per component to decrease with the decreasing LCC size because of both the embolisation of inlet and outlet vessels and the increased number of components. Importantly, all relationships between network measures depended on the intervessel connectivity structure and, for example, differences in connectivity structure between network components could lead to nonlinear relationships.

We evaluated the similarity of the behaviour of network properties between the physiological and SI spreading models in terms of the root mean square error (RMSE) between the time-dependent behaviour vectors. RMSE is defined for vectors of equal length. However, due to the different time scales of the two models, the SI model produced longer behaviour vectors than the physiological model (see Section 3.2 for details). We addressed this by interpolating more values for the physiological model vector with the numpy.interp function, and normalized the obtained RMSEs by the mean of the physiological behaviour vector for easier comparison.

### 2.5 Parameter estimation

The xylem network model of Mrad, Domec et al (2018) and Mrad, DM Johnson et al (2021) was parametrized for *Acer glabrum*. However, we selected *B. pendula* for the example species instead because this species shows a very broad geographic distribution pattern across the Eurasian continent. To re-parametrize the model for *B. pendula*, we used a set of parameters reported in the literature for large *B. pendula* branches (see Table 2).

To estimate the values of the probabilities 𝒩𝒫_*c*_, 𝒫_*c*_, 𝒫_*e,rad*_, and 𝒫_*e,tan*_, we constructed a set of xylem networks with the same size as those used for constructing the vulnerability curves so that there were 10 networks constructed with each combination of the probability parameters. We calculated for each network the vessel tissue fraction (VTF) (i.e. the fraction of the xylem cross-section area occupied by vessels) and vessel grouping index (GI) (i.e. the total number of vessels divided by the number of groups of adjacent vessels) and averaged them across the 10 networks per probability combinations. We then compared the observed VTF and GI values to the values reported for *B. pendula* (Fig. 2, see also Table 2). To this end, we assigned each parameter combination with two ranks, corresponding to VTF and GI, based on the absolute-valued difference between the value produced by the combination and the value reported in the literature. In other words, the parameter combination that produced the VTF value closest to the reference value from the literature, VTF_*ref*_, received rank 1, the combination that produced the second-closest value received rank 2, and so on. We then added together the VTF and GI ranks and selected the parameter combination corresponding to the lowest total rank. This optimization yielded 𝒩𝒫_*c*_ = 0.8, 𝒫_*c*_ = 0.9, 𝒫_*e,tan*_ = 0.9, and 𝒫_*e,rad*_ = 0.0. However, we did not want to completely exclude the possibility of radial VMs. Thus, we used 𝒫_*e,rad*_ = 0.01 instead of the optimization outcome. This parameter combination yielded VTF 0.26 and GI 2.61 (average across the 10 networks created for the selected probability parameter combination).

**Figure 2.**
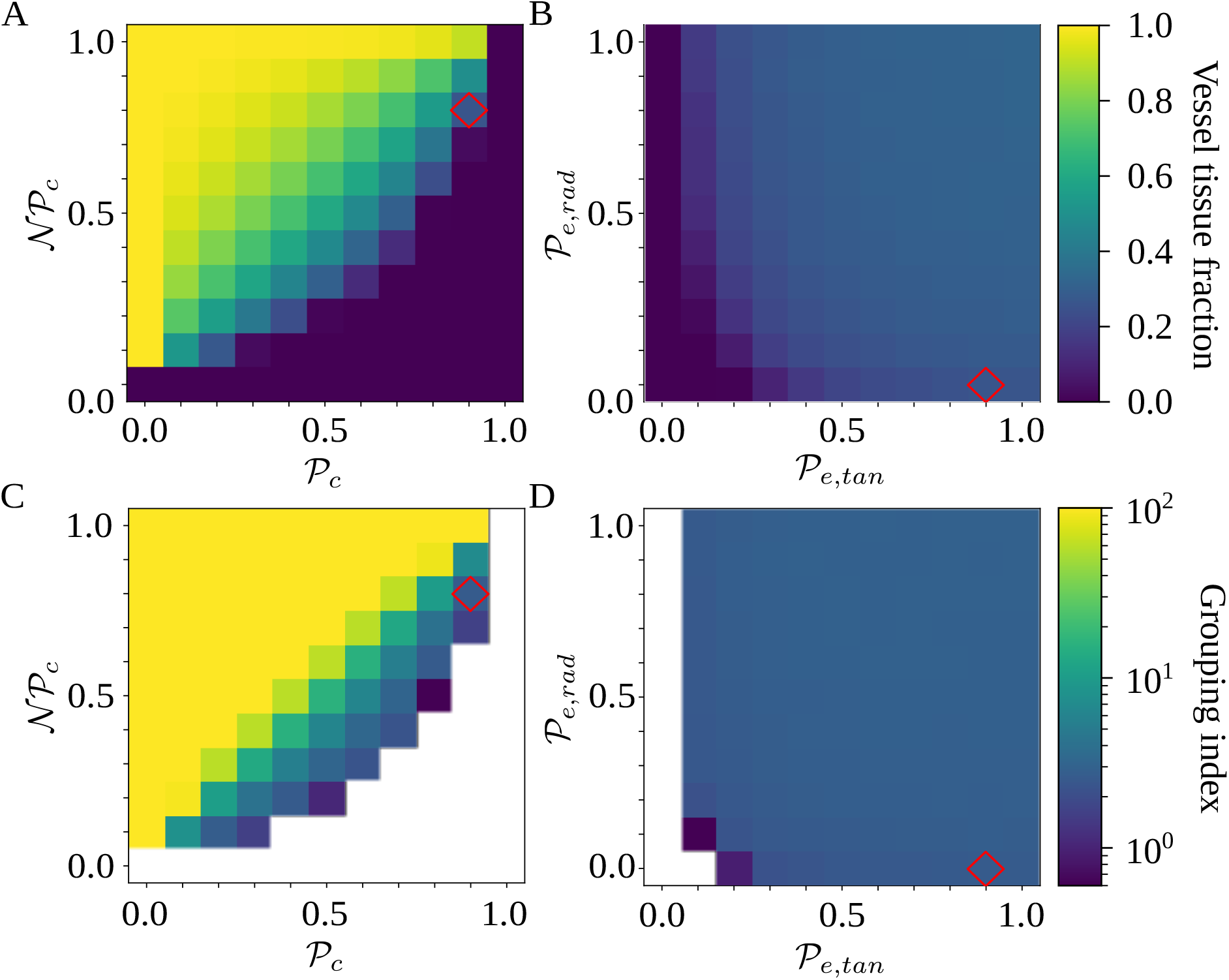
Optimization of probability parameters for xylem network construction. We constructed xylem vessel networks with different combinations of probabilities 𝒩𝒫_*c*_, 𝒫_*c*_, 𝒫_*e,rad*_, and 𝒫_*e,tan*_ and calculated the vessel tissue fraction (A, B) and grouping index (GI) (C, D) of each simulated network. Two 2D projections of the 4D space spanned by the four probability parameters are shown: for visualizing the effect of 𝒩𝒫_*c*_ and 𝒫_*c*_ (A, C), 𝒫_*e,rad*_ and 𝒫_*e,tan*_ are fixed to their optimal value, while the effect of 𝒩𝒫_*e,rad*_ and 𝒫_*e,tan*_ (B, D) is visualized fixing 𝒫_*c*_ and 𝒫_*c*_ to their optimal values. The red square shows the optimal parameter combinations. Note that the z scale of GI (C, D) is logarithmic; the blank cells indicate areas where GI = 0.

Network science commonly quantifies network structure in terms of average degree, defined as the average number of neighbours per network node, and network density, defined as

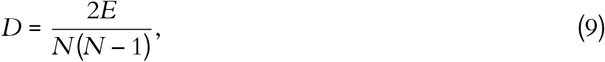

where *E* is the number of network links (here, number of VMs) and *N* is the number of nodes (here, number of vessels). These measures offer a basis for comparing the structure of networks of different sizes. The optimized probability parameters produced average degree 2.92 *±* 2.87E-2 and network density 8.56E-4 *±* 2.24E-5 (mean *±* across 100 networks constructed with the optimized probability parameters and used for simulating embolism spreading). This indicated that the simulated intervessel networks were sparse, similar to many real-world networks (e.g. Watts and Strogatz (1998) reported *D* = 2.71E–4 for a collaboration network of film actors, *D* = 5.41E–4 for the power grid of the western U.S. but notably more dense *D* = 4.98E–2 for the neuronal network of *Caenorhabditis elegans*). Further, the low standard deviation of the degree suggested that the VMs were relatively equally distributed across the vessels.

## 3. Results

### 3.1 SI model replicated physiological embolism spreading

To investigate if the stochastic SI model can approximate spreading of embolism in *B. pendula* xylem, we simulated embolism spreading with two models. Our physiological model was based on xylem anatomical and physiological characteristics as well as the network models of Loepfe et al (2007), Mrad, Domec et al (2018), and Mrad, DM Johnson et al (2021). The SI model, on the other hand, is a stochastic spreading model used to study epidemic spreading computationally in various fields, including network science (Hethcote 2000; Pastor-Satorras et al 2015). To compare the two models, we selected the free parameter of the SI model, the spreading probability β, separately for each *P* to minimize the difference in effective conductance *K*_*eff*_ between the two models. After this fitting, the SI model produced a vulnerability curve notably similar to the one produced by the physiological model (Fig. 3A), demonstrating that the spreading saturated at similar levels of embolism in both models. The optimized β value varied between 0 and 0.085, increasing almost monotonically as a function of *P* (Fig. 3B).

**Figure 3.**
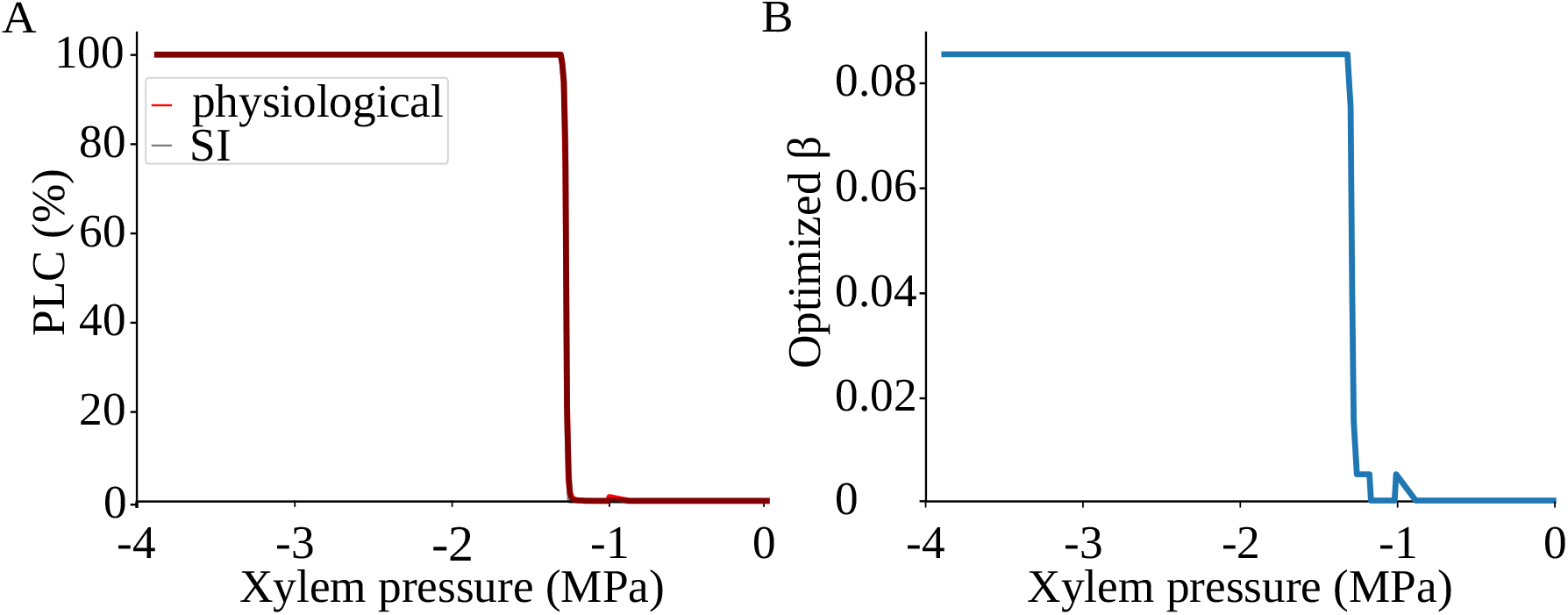
SI model replicates the vulnerability curve produced by the physiological model. A) PLC produced by the physiological (red) and SI (grey) models. For calculating the PLC, *K*_*eff*_ was averaged across 100 spreading simulations. The transition from normal xylem function to hydraulic failure happened almost step-like over a narrow *P* range; for further details, see the main text and section 4.3. B) The optimized SI spreading probability (β).

The shape of the vulnerability curves was sigmoidal, almost step-like, indicating a sharp transition between normal xylem function and hydraulic failure. The *P*_12_, *P*_50_, and *P*_88_ values (i.e. pressures where 12%, 50%, and 88% of *K*_*eff*_ were lost) of our physiological model were -1.26 MPa, -1.28 MPa, and -1.29 MPa, respectively. The full spreading process for *P*_12_, *P*_50_, and *P*_88_ is shown in Figs. 4 and 5.

**Figure 4.**
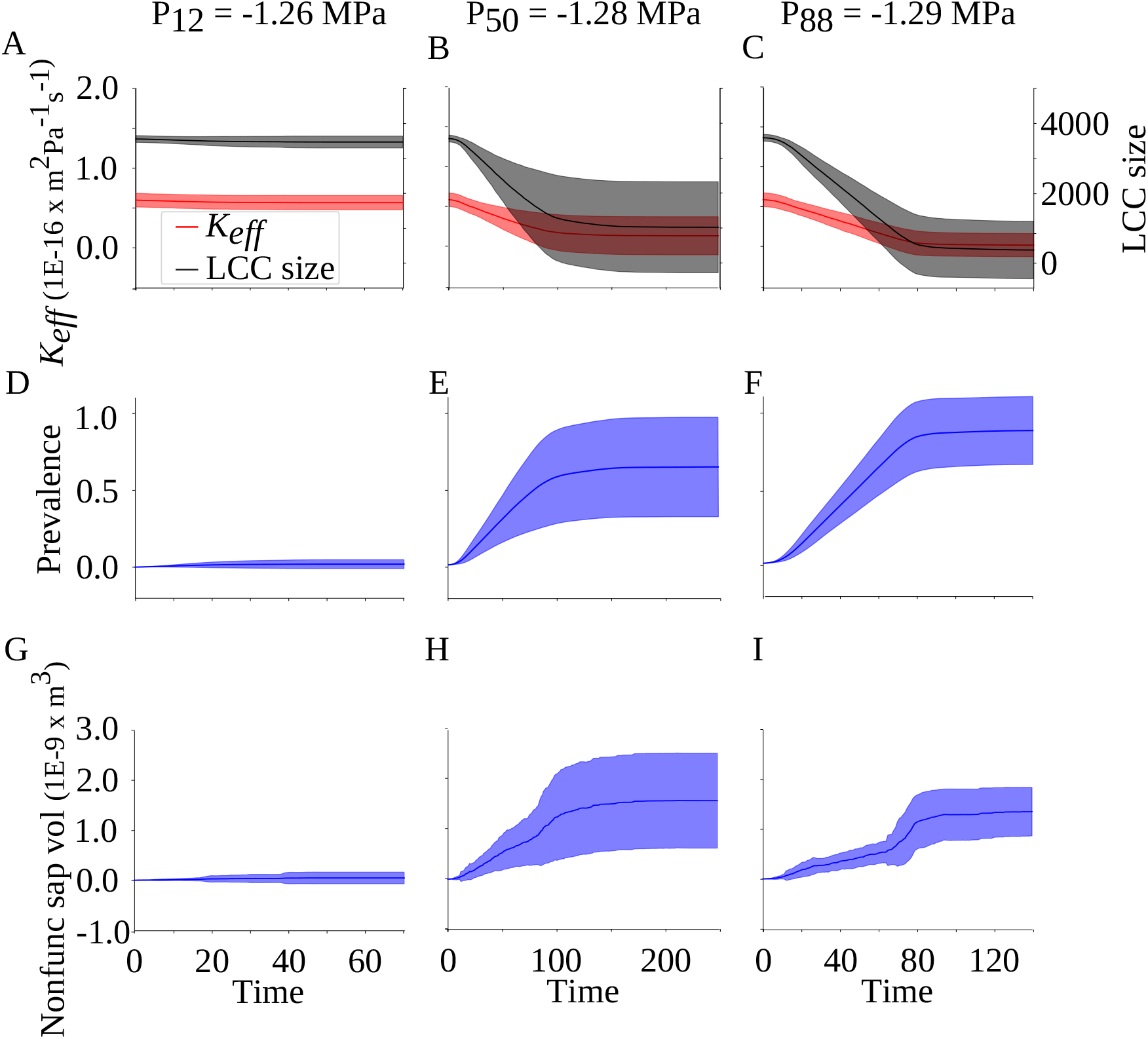
Behaviour of xylem network properties during embolism spreading simulated with the physiological model. *K*_*eff*_ and the largest connected component (LCC) size (A, B, C), prevalence (D, E, F), and nonfunctional sap volume (G, H, I) are monitored for *P*_12_ (A, D, G), *P*_50_ (B, E, H), and *P*_88_ (C, F, I) of the physiological spreading model. For each network property, the full line corresponds to the mean across 100 simulations, while the shadowed area shows the standard deviation across iterations. The x axis shows the dimensionless number of simulation time steps that does not directly map to absolute time. Note that the embolism spreading took different number of time steps to saturate at different *P* values, and the x axes thus differ between subplots.

**Figure 5.**
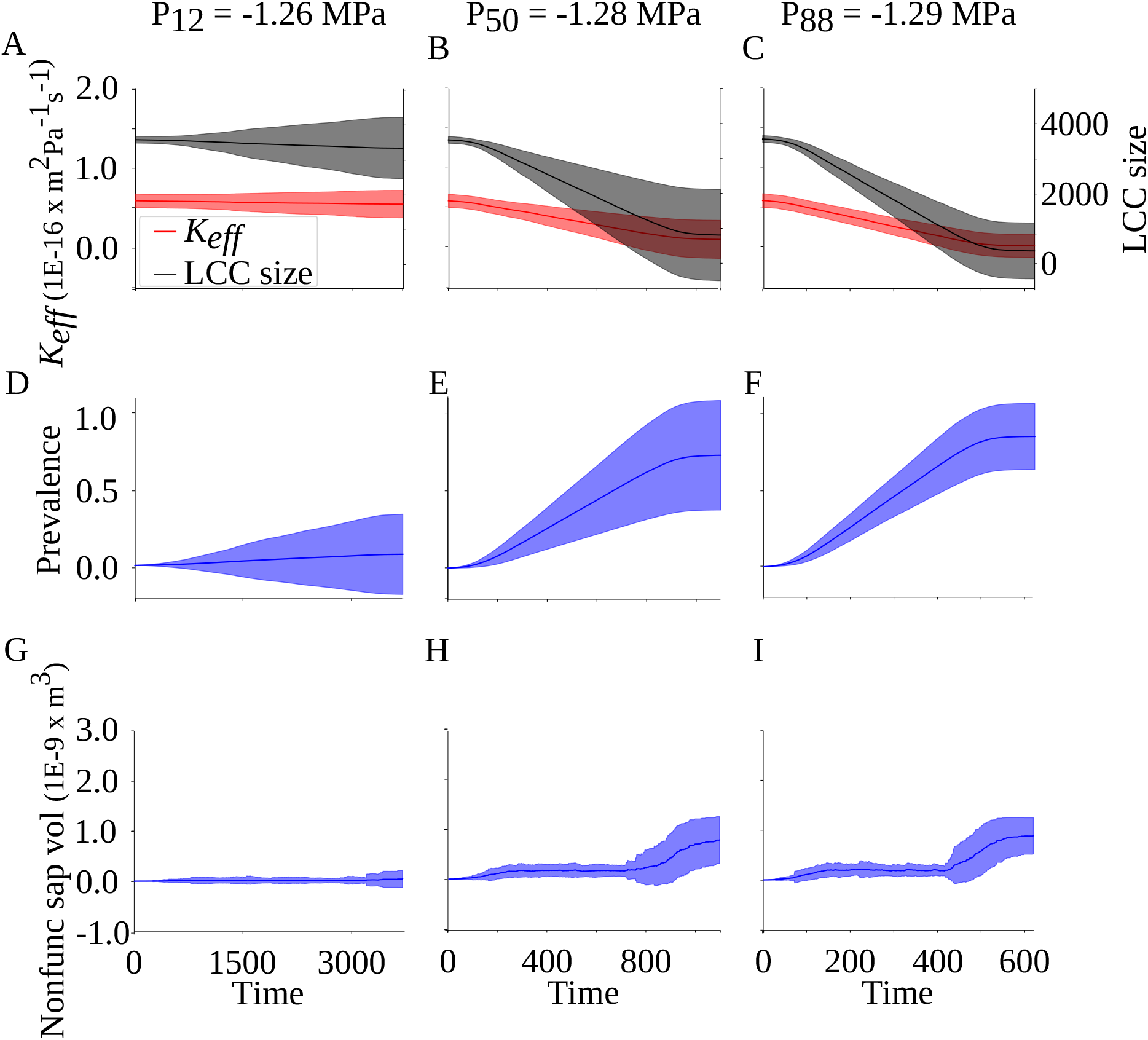
Behaviour of xylem network properties during embolism spreading simulated with the SI model. Similarly as in Fig. 4, *K*_*eff*_ and the largest connected component (LCC) size (A, B, C), prevalence (D, E, F), and nonfunctional sap volume (G, H, I) are monitored for *P*_12_ (A, D, G), *P*_50_ (B, E, H), and *P*_88_ (C, F, I), the full lines and shadowed areas show mean and standard deviation across simulations, and the x axis shows the number of simulation time steps. Similarly as in Fig. 4, note the difference in the x axis maximum values and spacing between subplots.

### 3.2 Network properties revealed the embolisation process

Vulnerability curves showed the final state of embolism spreading at each *P*. To explore how this final state was reached, we monitored several properties of the xylem network during the spreading process. In particular, we explored the behaviour of network properties as a function of increasing number of simulation time steps. At each time step, an embolised vessel could spread the embolism to all its sap-filled neighbours. In other words, the time step length corresponded to the time required by a sap-filled vessel to become fully embolised and able to spread the embolism further. This time depends on both xylem properties and external conditions, and therefore mapping simulation time steps to absolute time is not trivial.

In the case of the physiological spreading model (Figs. 4, S1A-B), the behaviour of *K*_*eff*_, the LCC size, and the functional LCC size at *P*_50_ and *P*_88_ was characterized by a quick decrease followed by a long, slower saturation phase, where the standard deviation across simulations increased. On the other hand, at *P*_12_, *K*_*eff*_, the LCC size, and the functional LCC size stayed stable over time. In the beginning of the simulation, the LCC was functional for all *P*s. However, for *P*_50_ and *P*_88_, the size of the largest nonfunctional component exceeded the size of the largest functional component in the beginning of the saturation phase (Fig. S1). Prevalence, or the fraction of embolised vessels, reflected the behaviour of *K*_*eff*_ : quick initial increase was followed by a long saturation phase at *P*_50_ and *P*_88_, while the behaviour at *P*_12_ was approximately static. Interestingly, at *P*_88_, the average final prevalence was lower than the final PLC, while at *P*_12_ and *P*_50_, the final prevalence and PLC were approximately equal. Behaviour of nonfunctional sap volume, defined as the total amount of water in vessels with no connection to either inlet, outlet, or both, supported the observation about the nonfunctionality of the LCC at more negative pressures: at *P*_50_ and *P*_88_, the nonfunctional sap volume increased as the embolism spreading proceeded and broke down network components. Finally, the average number of inlets and outlets showed the familiar pattern of quick initial decrease and slower saturation at *P*_50_ and *P*_88_ and static behaviour at *P*_12_ (Fig. S2). The number of outlets was higher than that of inlets because of the xylem network construction process: while a cell in the inlet row initiated a vessel with probability 𝒩𝒫_*c*_, all vessels that reached the outlet row ended automatically, thus yielding a higher number of outlet than inlet cells.

The most notable difference in the behaviour of network properties between the physiological model and the SI model (Figs. 5 and S1C-D) was the time scale. While the physiological spreading model saturated within less than 100 steps for *P*_12_, around 250 steps for *P*_50_, and around 140 steps for *P*_88_, the SI model necessitated over 3000 steps for *P*_12_, over 1000 steps for *P*_50_, and around 600 steps for *P*_88_. Despite the different time scales and the related rather high normalized RMSE values (Table 3), the behaviour of *K*_*eff*_, the LCC size, and the functional LCC size, prevalence, and the number of inlets and outlets per component were qualitatively similar between the two models, although the SI model yielded higher standard deviation at *P*_12_. The nonfunctional sap volume, instead, behaved differently in the SI model: it increased only in the later states of the simulation for both *P*_50_ and *P*_88_, and for *P*_50_, its final value remained lower than in the physiological model. However, for *P*_88_, the nonfunctional sap volume behaviour of the physiological model approached that of the SI model.

**Table 3.** RMSE between the network property behaviour vectors calculated with the physiological and SI spreading models.

| Property | RMSE |  |  |
| --- | --- | --- | --- |
| | $P_{12}$ | $P_{50}$ | $P_{88}$ |
| $K_{eff}$ | $2.41 \cdot 10^{-18}$, 0.0419<sup>a</sup> | $3.30 \cdot 10^{-17}$, 1.35 | $3.71 \cdot 10^{-17}$, 1.75 |
| LCC size | 68.7, 0.020 | 1920, 1.19 | 2190, 1.55 |
| Functional LCC size | 68.7, 0.020 | 2010, 1.30 | 2280, 1.69 |
| Prevalence | 0.015, 1.03 | 0.434, 0.980 | 0.551, 1.02 |
| Nonfunctional sap volume | $3.35 \cdot 10^{-11}$, 1.06 | $1.16 \cdot 10^{-9}$, 1.05 | $8.31 \cdot 10^{-9}$, 1.05 |

a RMSE and normalized RMSE; RMSE was normalized by the mean of the physiological behaviour vector.

### 3.3 SI model reproduced an empirical B. pendula vulnerability curve

So far, we have concentrated on how well the SI model replicated the outcome of the more detailed, physiological embolism spreading model. However, such physiological models are not available for all plant species due to limited availability of data on xylem physiology and anatomy. To demonstrate the potential of the SI model also in these cases, we next fitted the SI model against an empirical *B. pendula* vulnerability curve. Despite the difference in *P*_50_ between the physiological model and empirical data, the re-fitted SI model produced a vulnerability curve that closely matched the empirical one (Fig. 6). The optimized β varied between 0 and 0.11 and increased almost monotonically as a function of *P*, excluding small noise at the least negative *P* values and a peak in the beginning of the saturation zone. Both of these were probably artifacts related to the limited number of simulation iterations (100) used for optimizing the β value for each *P* value.

**Figure 6.**
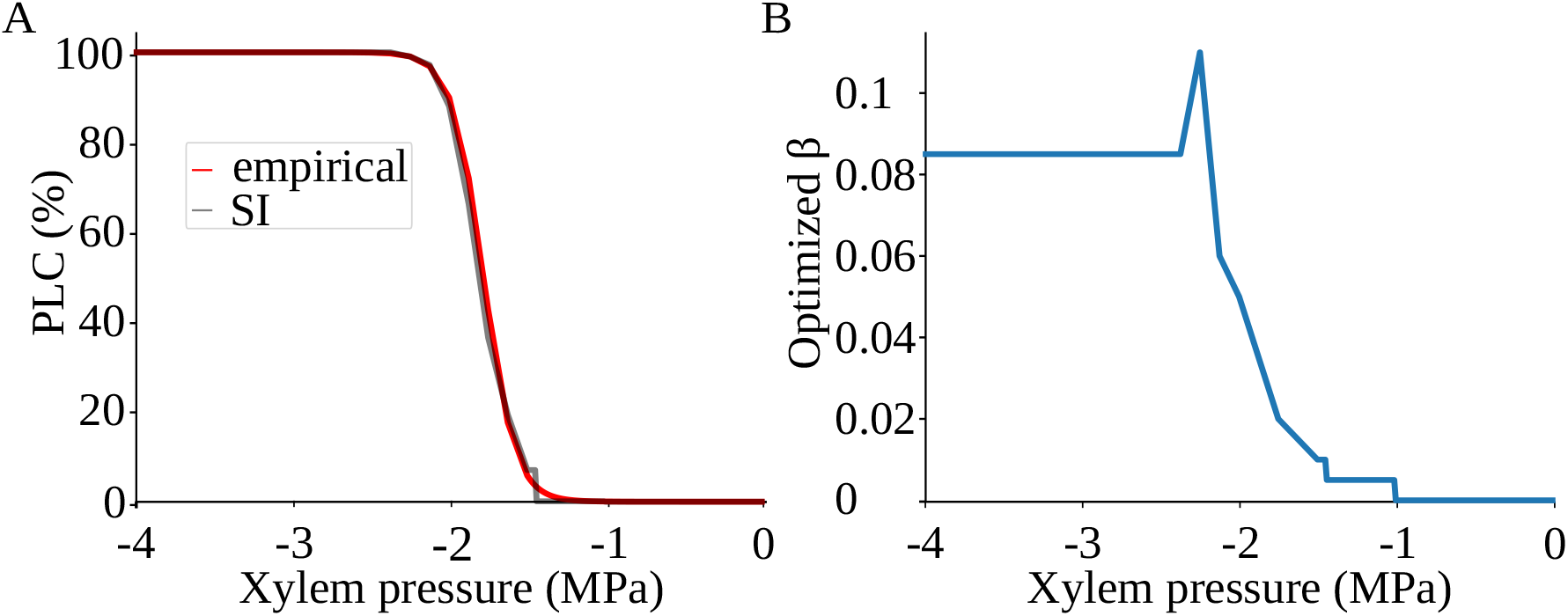
The SI model produces a vulnerability curve that matches empirical observations. A) The empirical vulnerability curve (red line) of *B. pendula* is from González-Muñoz et al (2018) and obtained as a sigmoidal fit to data. The modelled vulnerability curve (grey) is obtained by optimizing the SI spreading probability (β) for each *P*. B) The optimized β values used for constructing the SI vulnerability curve.

After re-fitting, the SI model allowed approximating the behaviour of network properties, for example prevalence, LCC size, and nonfunctional sap volume, during embolism spreading (Fig. S3). This behaviour is not directly measurable and cannot be addressed through the empirical vulnerability curve.

## 4. Discussion

### 4.1 SI model describes embolism spreading based on a single free parameter

In the present article, we have demonstrated that a stochastic model for embolism spreading in plant xylem, the SI model, produces similar results than previous, physiological embolism spreading models. Importantly, in contrast to physiological models that require detailed data on intervessel pit membrane function and anatomy, the SI model depends on a single spreading probability parameter that can be fitted against an empirical vulnerability curve. Therefore, the SI model opens new insights on embolism spreading without requiring detailed information about intervessel pit membrane structure.

The similarity between the physiological and SI models is not limited to the PLC measured at the final state of embolism spreading. Instead, the development of several xylem network properties across time was qualitatively similar in both models. While the earlier saturation in the physiological model and higher variance in the SI model yield rather high normalised RMSE values (see Table 3), the SI model is still able to qualitatively describe the dynamics of the xylem network structure during embolism spreading. While these dynamics can be empirically determined based on time-consuming computed tomography and image analysis (see, e.g., Choat, Badel et al (2016)), they are not accessible based on empirical vulnerability curves. Particularly the shape of the prevalence curve, fast initial growth followed by long saturation, is typical for the SI model and other compartment models without recovery (Pastor-Satorras et al 2015). Observing this shape of prevalence curve for both the physiological and SI models strongly supports modelling embolism spreading with the SI model.

The most notable difference between the SI and physiological models is the time scale of the saturation of embolism spreading: the physiological model requires around 10-fold less time to saturate than the SI model. However, mapping the simulation time steps to absolute time is not straightforward. Both spreading models consider embolisation of a vessel immediate so that all vessels embolised at time *t* can spread embolism further at time *t* + 1. However, this is a simplification: in reality, the time required for a vessel to become fully embolised and able to spread the embolism to its neighbours is not constant but varies depending on, for example, the sap pressure, status of the neighbouring vessels, and sap lipid concentration. Therefore, we consider the ability of the SI model to reproduce the outcomes of the physiological model qualitatively more important than the difference in the simulation timescales. In particular, we do not recommend using either of the spreading models to estimate the absolute time scale of embolism propagation.

Further, it is important to understand the different meaning of saturation in the two models. In both models, embolism spreading slows down when the number of susceptible vessels decreases, as described above. In the deterministic physiological model, this leads to true saturation: at some point, all vessels that can be embolised at a given *P* are embolised and the spreading stops. The SI model, on the other hand, would, due to its stochasticity, yield 100% prevalence in an infinitely long simulation. Therefore, at the saturation point, the spreading does not stop but becomes extremely slow.

In our simulations, already relatively low spreading probabilities (around 0.2) yield very high prevalences at the saturation point. Therefore, fitting the model for *P* values where the full network does not become embolised has yielded very low spreading probability (β) values (0.01, 0.04, and 0.075 for *P*_12_, *P*_50_, and *P*_88_, respectively) associated with long saturation times. Larger βs would guarantee faster saturation but also produce undesirably low final *K*_*eff*_. Another way to shorten the runtime of the SI model would be to record the number of simulation steps the physiological model needed to saturate, *n*_*sat*_, and optimize β based on the *K*_*eff*,*SI*_ (*n*_*sat*_) yielded by the SI model after *n*_*sat*_ steps. However, while this strategy would make the simulations slightly faster, it would weaken the correspondence of the behaviour of network properties during the spreading process between the two models. Further, this optimization strategy cannot be used when optimizing the spreading probability against an empirical vulnerability curve as the empirical saturation time is typically unknown.

### 4.2 Xylem network breakdown contributes to hydraulic failure

In both the physiological and the SI model, the largest connected component of the xylem network becomes nonfunctional, that is, remains sap-filled but loses its connection to network inlet or outlet and thus stops contributing to sap flow, during embolism spreading. Further, for more negative *P*s, the final prevalence, or fraction of embolised vessels, is lower than the PLC. In other words, especially at more negative pressures, hydraulic conductivity is lost faster than the xylem sap volume becomes reduced. These results indicate that while our hypothesis about the network breakdown as the main cause of hydraulic failure holds only for the more negative *P*s, the disruption of the xylem network structure that cuts down the sap pathways is an equally important reason for hydraulic failure as the drying up of the xylem at all pressure values. This observation is supported by earlier results on angiosperms, where leaf water potential and transpiration reduce faster than the fraction of sap-filled vessels in stem xylem (Gauthey et al 2022), and conifers, where xylem pressures close to *P*_50_ cause 25% loss in branch tissue water content (Rosner et al 2021). Our observations on the nonfunctional sap volume confirm this result: significant amounts of sap remain in the xylem tissue even when 88% of conductance is lost.

While the final nonfunctional sap volume indicates presence of sap in the xylem also after hydraulic failure in both models, the behaviour of nonfunctional sap volume during the spreading process differs between the models. In the physiological model, we observe a nearly monotonic, continuous increase, while in the SI model, the nonfunctional sap volume is initially low and increases only close to the saturation point. This difference reflects the deterministic and stochastic natures of the two models. Let *C* be a part of the xylem that is connected to the rest of the network, and thus to inlet and outlet, only through vessel *I*. Now, embolisation of *I* cuts *C* apart from the network, and it becomes a nonfunctional component. In the physiological model, embolism cannot further spread to *C* if the BPP between *I* and its neighbours in *C* is larger than *P*. Therefore, the total volume of vessels in *C* contributes to increase of the nonfunctional sap volume until the end of the simulation. In the SI model, on the other hand, embolism can still spread from *I* to *C* with probability β, and the nonfunctional sap volume decreases as vessels in *C* become embolised. At the level of the whole network, the simultaneous addition of new nonfunctional components and removal of embolised vessels in existing nonfunctional components keeps the overall nonfunctional sap volume lower than in the physiological model. At very high *P* values, BPP values larger than *P* become rare and embolism can thus spread to most of the nonfunctional components. Indeed, at high *P*s, the behaviour of nonfunctional sap volume in the physiological model approaches that observed in the SI model.

### 4.3 Network structure and model parameters affect modelled embolism spreading

Historically, intervessel connectivity has been seen as a safety mechanism, following the reasoning of Carlquist (1977): in presence of numerous, parallel vessels, an embolism in a single vessel is less probable to cause hydraulic failure in the entire xylem. However, under certain conditions tight intervessel connectivity can rather increase than decrease vulnerability of the xylem (Ewers et al 2007; Loepfe et al 2007; Wheeler et al 2005). From the viewpoint of pit membrane structure, this finding is in line with the pit area hypothesis (i.e. large pit membrane areas are more vulnerable to embolism spreading than small areas) but not with the rare pit hypothesis (i.e. embolism resistance depends on a single, exceptionally large pit membrane pore) (Kaack, Weber et al 2021; Wheeler et al 2005). In general, this finding indicates a more complex relationship between intervessel networks, sap flow, and embolism spreading in xylem than assumed before. Meanwhile, modern advanced imaging techniques, such as computed tomography (Brodersen et al 2011; Choat, Badel et al 2016), have yielded new understanding of the intricacies of xylem architecture. Embolism propagation is a dynamic process that takes place on top of the intervessel network. Thus it is clear that appropriate representation of the complex 3D structure of the xylem will be critical to accurately simulate embolism propagation.

The behaviour of prevalence as a function of time depends on network structure and average degree of network nodes (Barthélemy et al 2005; Pastor-Satorras et al 2015; Vespignani 2012). In real-world networks, also network size affects the prevalence. In the present article, we used relatively small networks (100 × 10 × 100 potential vessel elements). We did not investigate the effects of the simulated xylem network size on *P*_50_. Therefore, applying the model on networks of a different size should be considered with care.

In the case of physiological models, the final state of embolism spreading is affected both by the underlying xylem network structure and the modelling of intervessel pit membrane function. In our physiological model, we modelled the intervessel pit membrane as a 3D object instead of the more traditional 2D surface model. Importantly, in the 3D model, intervessel pit membrane pores are not 2D openings but channels that comprise multiple pore constrictions. This affects the estimation of BPP across the pit membrane: while most 2D models, including the one used by Mrad, Domec et al (2018) and Mrad, DM Johnson et al (2021), build on the rare pit hypothesis, simulations using the 3D models show that the probability to find exceptionally large pit membrane pores is very low especially in species with thicker pit membranes (Kaack, Weber et al 2021). Therefore, the 2D models may produce artificially high embolism propagation probabilities and overestimate species’ vulnerability to hydraulic failure due to embolism. The intervessel pit membrane of *B. pendula* is relatively thin (205 nm according to Kaack, Weber et al (2021)). Therefore, the effect of the pit membrane model selection may not be particularly large in our analysis. However, the difference between the 2D and 3D pit models becomes more prominent when the pit membrane thickness increases. Therefore, it is a notable source of error for many angiosperms (Kaack, Weber et al 2021). Empirical observations on the movement of nanoparticles through pit membranes support the selection of the 3D model (Kaack, Weber et al 2021; Zhang et al 2024).

The *P*_50_ value of our physiological model, -1.28 MPa, was slightly less negative than the empirically measured values for *B. pendula*, which varied from -1.59 to -1.9 for six vulnerability curves fitted based on the flow-centrifuge measurements (Dulamsuren et al 2019; González-Muñoz et al 2018). We consider these values largely comparable because a 10% to 30% error from the true *P*_12_, *P*_50_, and *P*_88_ values is acceptable and equal to the typical disagreement in embolism resistance measured by different methods on the same species (Silva, Pereira et al 2024; Silva, Pfaff et al 2024; Yang et al 2023). However, there are also potential error sources that may explain the difference between the modelled and empirical *P*_50_ values. First, the embolism resistance measured by González-Muñoz et al (2018) could have been slightly overestimated due to time-based shifts in vulnerability curves generated with a flow-centrifuge (Silva, Pfaff et al 2024) and the temporal buffering effect of water release from embolism on water potential declines and embolism spreading (Hölttä et al 2009; Vergeynst et al 2015). However, the potential buffering effect is minor and needs to be investigated in further detail.

On the other hand, the difference between the modelled and empirical *P*_50_ values may relate also to the pit membrane model, which yielded for single pit membranes BPP values slightly higher than the empirical *P*_50_ values of the same species, as suggested also by Kaack, Weber et al (2021). This indicates that despite its more realistic view of the pit membrane as a 3D object, the model still describes some interactions between liquids, surfactants, and gas nanobubbles at the pit membrane inaccurately. Understanding these interactions is an important direction for future research.

Further, our physiological model selects the seed vessels for embolism spreading in a way that is not entirely realistic. First, a certain amount of embolised vessels is known to exist in intact plants at native pressures (Torres-Ruiz et al 2016), while understanding the origin of these initial embolisms remains a subject for further research (Carmesin et al 2023). Therefore, selecting a single seed per network component may be too little. Second, while we select the seed vessels at random, in reality these vessels are more common in the most central parts of the xylem (Choat, Badel et al 2016). However, underestimating the number of seed vessels cannot explain the observed difference between the empirical and observed values, since it should make the modelled *P*_50_ value more, not less, negative. Further, the effect of ignoring the distribution of seed vessel locations is probably negligible, as in our model the properties of seed vessels, most importantly their degree (i.e. number of neighbours) that affects the SI spreading process, are independent of the location in the xylem.

In addition to the difference between the *P*_50_ values, the output of our physiological model differed from empirical data in terms of the vulnerability curve shape. Our model produced *P*_12_ and *P*_88_ values closer to each other than observed empirically (-1.26 MPa and -1.29 MPa vs. -1.62 MPa and -2.00 MPa in González-Muñoz et al (2018)). Thus, in our simulations PLC increased as a function of *P* faster than what has been observed empirically, and the transition from normal xylem function to hydraulic failure happened over an unrealistically narrow range of pressures. Since our investigated pressure range was particularly dense around the transition point (for details, see section 2.3.3), the steep increase in PLC is a feature of the physiological model itself. Most probably, this discrepancy is related to the fact that the model considers embolism events immediate so that a vessel can spread embolism further right after becoming embolised. In reality, embolism spreads from a newly embolised vessel only after a certain gas concentration is reached (Pereira et al 2023; Silva, Bujnowski et al 2025), and ignoring this delay may make the vulnerability curve more step-like. In future, this problem can be addressed by adding to the model a stochastic waiting time, during which a newly embolised vessel cannot yet spread embolism, and fine-tuning the related compartment model (for details, see section 4.4).

Finally, our model concentrated on embolism spreading in a network formed by a single type of conduits, vessels. In *B. pendula*, imperforate tracheary elements (ITEs) are unlikely to be conductive (as Sano et al (2011) observed for a related species, *B. japonica*). However, they may still affect the resistance against embolism propagation through their mechanistic support role, and excluding them from the model is thus a potential, although minor, source of error. Further, including ITEs in the model would yield a more realistic picture of embolism spreading for the many angiosperm species with ITEs as a second type of conduits in addition to vessels (Esteban et al 2024; Olson 2023). To this end, a promising approach would be a multilayer network model (Kivelä et al 2014), where networks of different conducting cells, including vessels and ITEs, would form layers of their own while the interconnectivity between conduits of different types would be represented by the interlayer links.

### 4.4 Which compartment model is best suited for modelling embolism spreading?

The SI model belongs to the family of compartment models. These models divide individuals (here, xylem vessels) into several groups, or compartments, and monitor the transitions between these groups (Hethcote 2000; Pastor-Satorras et al 2015). As the simplest member of the family, the SI model comprises only two compartments: the susceptible (S) (i.e. non-embolised) and infected (I) (i.e. embolised) individuals. More complicated compartment models include, for example, recovery from infection by returning to the susceptible compartment (the SIS model) or removal of infected individuals through immunity or death (the SIR model) (Pastor-Satorras et al 2015).

The SI model is a strong simplification in the case of many infectious diseases (Pastor-Satorras et al 2015). However, it describes embolism spreading more naturalistically than other compartment models. For example, the SIS model would suggest recovery from embolism through refilling of embolised vessels. However, such refilling is rarely observed in woody plants under negative pressure (Choat, Brodersen et al 2015; Choat, Nolf et al 2019; Gauthey et al 2022; Knipfer et al 2015), and even in non-woody plants these mechanisms take place at time scales longer than those considered in our spreading simulations (Stewart et al 2025). In *B. pendula*, refilling takes place under seasonal positive pressures, but it remains unclear if this is possible under normal, negative xylem pressures (Choat, Brodribb et al 2018).

On the other hand, as discussed in section 4.3, a newly-embolised vessel can spread embolism further only after a waiting time. Therefore, the model could be further improved by adding an exposed (E) compartment for vessels that are already embolised but do not yet spread the embolism. In this hypothetical SEI model, the E-I transition could happen at a constant rate based on empirical observations. On the other hand, if the S-E transition was defined as a snap-off event yielding to an increased number of gas nanobubbles in the exposed vessel, the E-I transition could be also stochastic and defined by the probability of nanobubble expansion (Ingram et al 2024). Another potential extension of the SI model would be modelling several, possibly interacting, spreading processes on the same xylem network. Such model could describe, for example, the simultaneous spreading of embolism and fungi in the xylem.

Earlier, Roth-Nebelsick (2019) has modelled embolism spreading with the SIR model, where the removed (R) compartment consisted of areas that can no more spread embolism due to lack of susceptible neighbours. Torres-Ruiz et al (2016), on the other hand, has modelled embolism spreading between tracheids using an SI model variant that allowed both spontaneous embolism and spreading between orthogonally adjacent tracheids. While these models were able to reproduce empirical results, they considered the xylem as a continuous medium, ignoring the intervessel connectivity. However, the connectivity structure plays an important role in xylem hydraulics (Avila, Kane et al 2023; Cai et al 2014; Guan et al 2021; Loepfe et al 2007) and in spreading processes in general (S Johnson 2024; Newman 2002; Pastor-Satorras et al 2015). Our approach, running the SI model on top of simulated xylem networks, combines the small number of free parameters typical for compartment models with the possibility to investigate how intervessel connectivity affects embolism spreading and to observe the changes in the xylem network structure during the spreading process.

### 4.5 Embolism spreading is a directed percolation phenomenon

In addition to exploring the effects of intervessel connectivity on xylem hydraulics, running spreading models on top of the intervessel network allows investigating embolism spreading with tools adopted from network science and applied to investigate a diverse set of real-world systems (Newman 2003). From this viewpoint, embolism spreading is an instance of a well-known concept of theoretical physics, directed (bond) percolation, that is widely applied to study inherently directed phenomena such as filtering, or percolation, of fluids through a porous media (Badie-Modiri et al 2022; Hinrichsen 2000). In network science, a theoretical mapping has been established between directed percolation and SI spreading processes on top of networks (Badie-Modiri et al 2022; Kenah and Robins 2007; Newman 2002; Pastor-Satorras et al 2015).

Directed percolation processes are characterized by a sudden transition between permeable and impermeable phases (Badie-Modiri et al 2022; Hinrichsen 2000). These transitions are observed in terms of two parameters: as the value of the free control parameter changes, the system transitions between regions where an observable order parameter gets either zero or non-zero values (Badie-Modiri et al 2022; Hinrichsen 2000). In network science, the two phases correspond to connected and fragmented network structures, and the percolation transition is observed when a network breaks down due to link removal (Newman 2003). Here, the control parameter is the fraction of removed links, while the most commonly observed order parameter is the size of the giant component, defined as a component that spans a finite fraction of the network when the network size grows to infinity. In empirical studies, the size of the largest connected component approximates the giant component size. As predicted by the percolation theory, most networks are relatively robust against link removal until a critical fraction of links is achieved. After this, the network quickly breaks apart, as indicated by a sudden drop in the largest connected component size. The exact value of the critical fraction of removed links, known as the percolation threshold, depends on the degree distribution of the network (Newman 2003).

When considering the SI spreading process as directed percolation, the control and order parameters correspond to the spreading probability and fraction of infected nodes, correspondingly. The percolation is directed because of the causal nature of infection spreading: while the infection can spread along network links in all spatial directions, the spreading along an additional time axis is limited, since nodes can infect their neighbours only after they themselves have been infected. Because of the lack of recovery mechanisms in the SI model, any non-zero spreading probability will eventually lead to a fully infected network. Therefore, the two states, between which the transition takes place, are not defined by the final prevalence but the prevalence at the saturation phase, where spreading slows down significantly (see Figs. 4 and 5). If this saturation happens only when prevalence is approximately 100%, the entire network will become infected in a finite time also when the network size grows to infinity, leading to non-zero order parameter values. On the other hand, if the saturation happens at a lower prevalence value, the time required to infect the entire network grows to infinity together with network size.

In embolism spreading, we see the sigmoidal shape characteristic for the percolation transition in the vulnerability curves that visualize the PLC as a function of *P* (see Fig. 3A and, e.g., Avila, Guan et al (2022) and González-Muñoz et al (2018)). This is not surprising, as *K*_*eff*_ depends on the percolation order parameter, the largest connected component size. This shape indicates that the plant xylem is relatively resistant to embolism until a certain threshold is reached, after which the embolism quickly spans the whole vessel network, leading to a percolation-like collapse of network structure (Avila, Guan et al 2022; Delzon and Cochard 2014). There are two explanations for the sigmoidal shape of the vulnerability curve. First, in all spreading processes, the spreading speed increases with the number of infected individuals since, in the early stages, the number of susceptible individuals (here, sap-filled vessels) with infected neighbours increases faster than the number of infected individuals (here, embolised vessels) itself (Pastor-Satorras et al 2015). Second, in plants specifically, above the embolism formation threshold, embolism propagation depends less strictly on the pressure difference across intervessel pit membranes, possibly because of the increased role of gas diffusion in embolism spreading at more negative *P* values (Avila, Guan et al 2022; Silva, Pereira et al 2024). This decreases the effect of pit safety mechanisms and thus makes the xylem more vulnerable to embolism spreading (Avila, Guan et al 2022).

The role of gas diffusion in speeding up embolism spreading is two-fold. On one hand, after becoming embolised, a vessel needs to reach the atmospheric gas concentration before spreading embolism to neighbouring vessels (Pereira et al 2023; Silva, Bujnowski et al 2025). This typically happens through diffusion both radially and from neighbouring sap-filled vessels, which, as discussed in section 4.3, may act as a temporary buffer mechanism against embolism in the neighbouring vessels (Silva, Bujnowski et al 2025). However, when the fraction of embolised vessels exceeds the percolation threshold, the high availability of air in the xylem speeds up diffusion to newly-embolised vessels, thus weakening this buffer mechanism (Avila, Kane et al 2023; Silva, Bujnowski et al 2025; Silva, Pereira et al 2024). On the other hand, after the embolised vessels have reached the atmospheric gas concentration, they become air sources themselves and can spread embolism both through snap-off events and diffusion (Silva, Pfaff et al 2024). Since the detailed mechanisms related to diffusion in embolism spreading are still not fully known, they are not included in our physiological model. However, fitting the SI model against empirical data can account for the increased role of diffusion through an increased spreading probability, β, above the embolism formation threshold. Interestingly, central evidence for the role of diffusion comes from the increase of PLC as a function of time when *P* is kept constant. While the SI model does not explicitly model diffusion, similar increase of PLC over time is seen due to the stochastic nature of the model.

Defining the exact mapping between embolism spreading and directed percolation remains as a subject for future research. However, our results on the ability of the SI model to describe embolism spreading in xylem, together with the known theoretical equivalence between SI spreading and directed percolation, demonstrate that embolism spreading belongs to the universality class of directed percolation. Mrad, DM Johnson et al (2021) has already proposed applying percolation theory to understand the connection between the average connectivity between vessels and embolism threshold. Finding the exact mapping to directed percolation will open further application possibilities for percolation theory, including the analytical and numerical results on the size distribution of embolised components and behaviour of prevalence across time.

## 5. Conclusion

We have demonstrated that a stochastic model for embolism spreading, the SI model, produces vulnerability curves comparable to those produced by a physiological model or extracted from empirical data. The succesful fitting of the SI model against the empirical vulnerability curve of *B. pendula* shows that the model can be used to investigate embolism spreading dynamics also in species, for which detailed data on xylem physiology are not available. Running spreading models on top of a simulated xylem network allows exploring the effects of intervessel connectivity on embolism spreading and monitoring xylem network properties during the spreading process. In particular, we observed that during the spreading, the largest connected component of the intervessel network becomes nonfunctional and does not further contribute to sap flow. This indicates that conductance loss is caused by both direct embolism and groups of non-embolised vessels being cut off from the functional xylem network. Combining the SI spreading model with intervessel connectivity structure also sets embolism spreading into a broader context of network phenomena as an instance of directed percolation.

## Supplementary information

### Supplementary figures

**Fig. S1:**
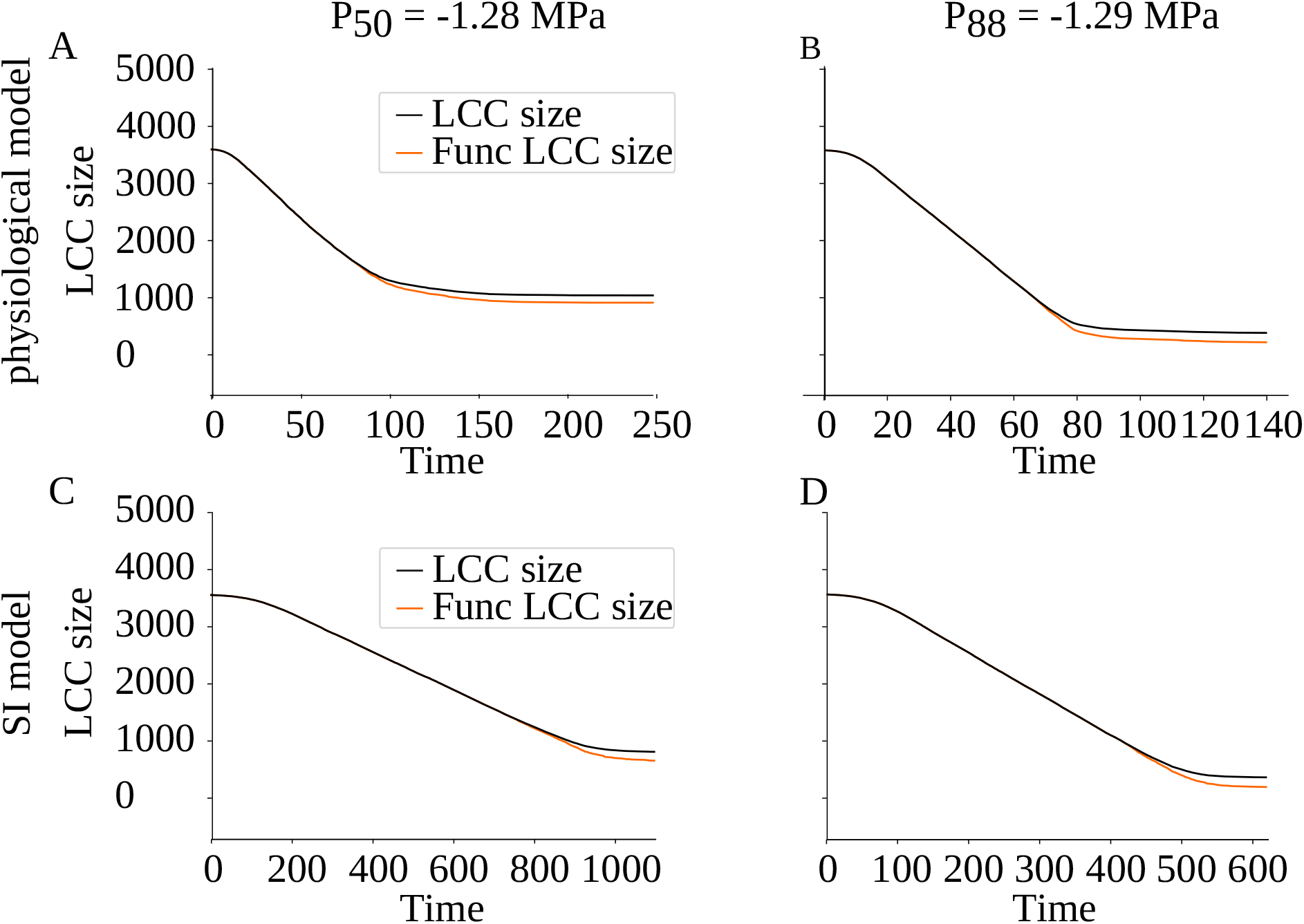
Largest connected component became nonfunctional at more negative xylem pressures. Behaviour of the LCC size (black) and functional LCC size (cyan) as a function of simulation time for the physiological (A, B) and SI (C, D) models at *P*_50_ (A, C) and *P*_88_ (B, D). At *P*_12_, LCC was always functional.

**Fig. S2:**
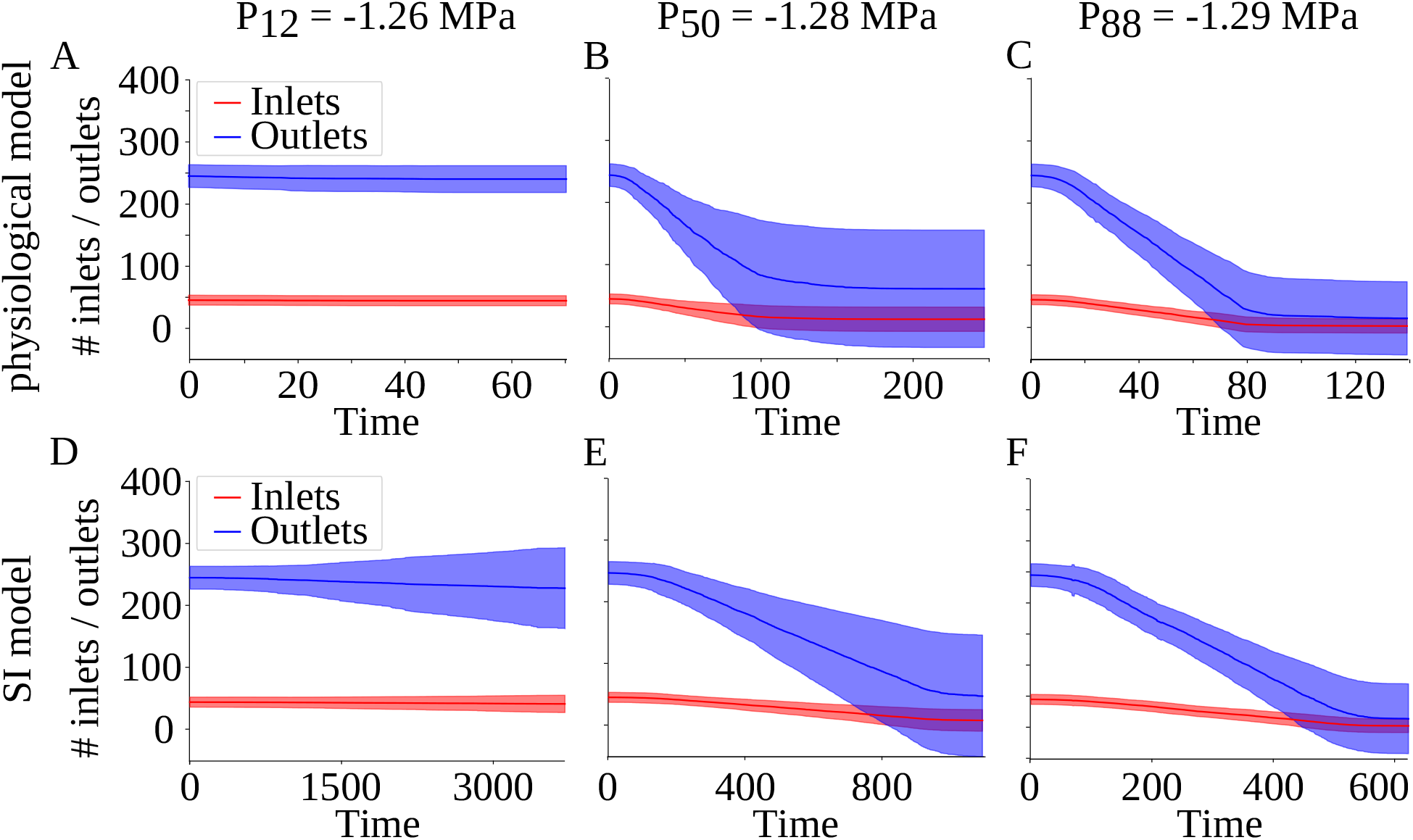
Behaviour of the average number of inlet and outlet vessels per component as a function of simulation time steps for the physiological (A, B, C) and SI (D, E, F) models at *P*_12_ (A, D), *P*_50_ (B, E), and *P*_88_ (C, F).

**Fig. S3:**
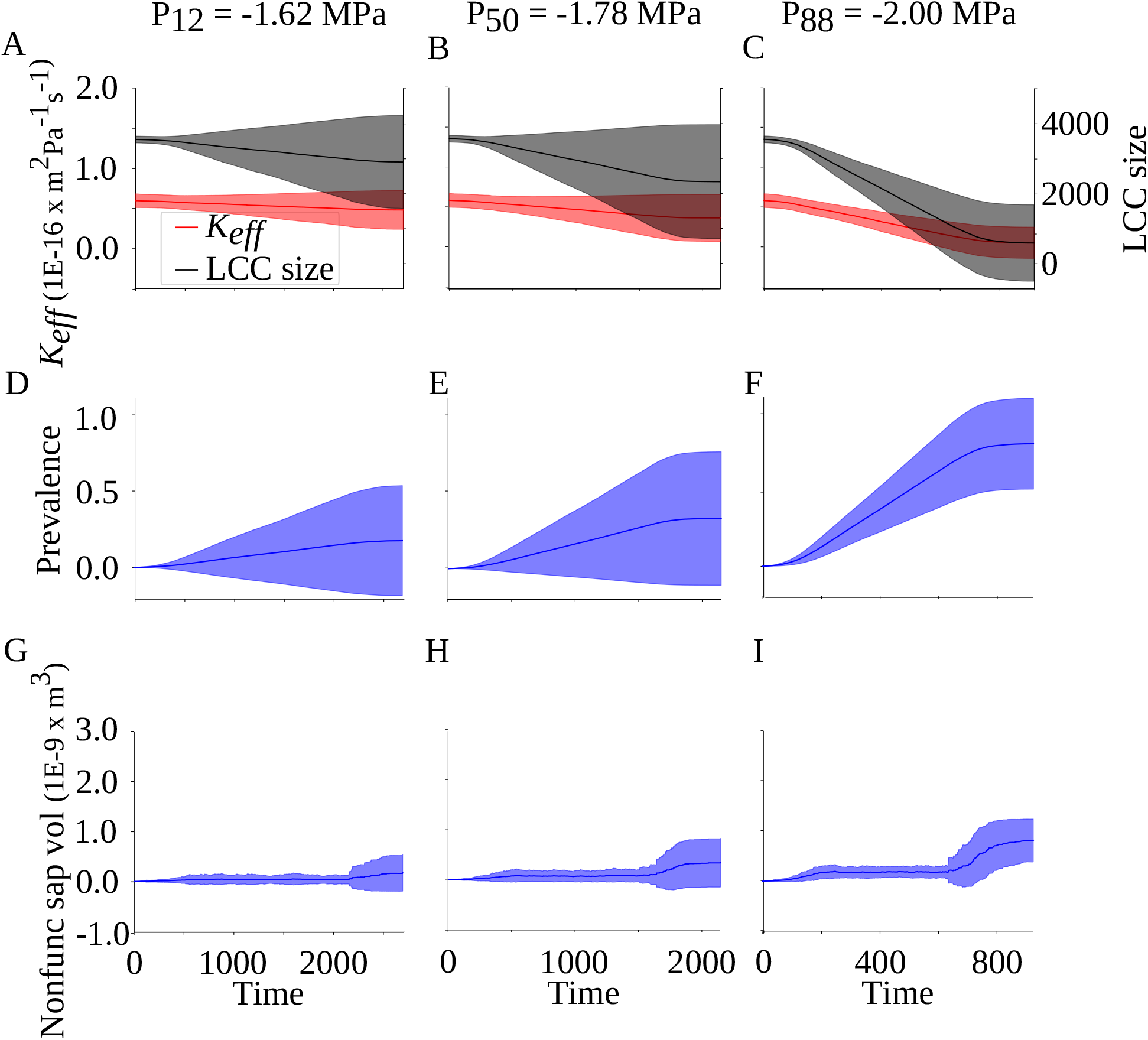
The SI spreading model fitted to empirical data allowed estimating the behaviour of network properties during embolism spreading based on an empirical vulnerability curve. *K*_*eff*_ and the largest connected component (LCC) size (A, B, C), prevalence (D, E, F), and nonfunctional sap volume (G, H, I) are monitored for *P*_12_ (A, D, G), *P*_50_ (B, E, H), and *P*_88_ (C, F, I). The full lines and shadowed areas show mean and standard deviation across simulations, and the x axis shows the number of simulation time steps.

## References

Adams HD, Zeppel MJ, Anderegg WR, Hartmann H, Landhäusser SM, Tissue DT, Huxman TE, Hudson PJ, Franz TE, Allen CD et al (2017) A multi-species synthesis of physiological mechanisms in drought-induced tree mortality. Nature Ecology & Evolution 1(9), 1285–1291.

Alber M, Petit G and Sellin A (2019) Does elevated air humidity modify hydraulically relevant anatomical traits of wood in Betula pendula? Trees 33, 1361–1371.

Anderegg WR, Klein T, Bartlett M, Sack L, Pellegrini AF, Choat B and Jansen S (2016) Meta-analysis reveals that hydraulic traits explain cross-species patterns of drought-induced tree mortality across the globe. Proceedings of the National Academy of Sciences 113(18), 5024–5029.

Avila RT, Guan X, Kane CN, Cardoso AA, Batz TA, DaMatta FM, Jansen S and McAdam SA (2022) Xylem embolism spread is largely prevented by interconduit pit membranes until the majority of conduits are gas-filled. Plant, Cell & Environment 45(4), 1204–1215.

Avila RT, Kane CN, Batz TA, Trabi C, Damatta FM, Jansen S and McAdam SA (2023) The relative area of vessels in xylem correlates with stem embolism resistance within and between genera. Tree Physiology 43(1), 75–87.

Badie-Modiri A, Rizi AK, Karsai M and Kivelä M (2022) Directed percolation in random temporal network models with heterogeneities. Physical Review E 105(5), 054313.

Barrat A, Cattuto C, Kivelä M, Lehmann S and Saramäki J (2021) Effect of manual and digital contact tracing on COVID-19 outbreaks: a study on empirical contact data. Journal of the Royal Society Interface 18(178), 20201000.

Barthélemy M, Barrat A, Pastor-Satorras R and Vespignani A (2005) Dynamical patterns of epidemic outbreaks in complex heterogeneous networks. Journal of Theoretical Biology 235(2), 275–288.

Bhat K and Kärkkäinen M (1981) Variation in structure and selected properties of Finnish birch wood. IV. Silva Fennica 15(1).

Bosshard HH and Kučera L (1973) Die dreidimensionale Strukturanalyse des Holzes—Erste Mitteilung: Die Vernetzung des Gefäßsystems in Fagus sylvatica L. European Journal of Wood and Wood Products 31(11), 437–445.

Brodersen CR, Lee EF, Choat B, Jansen S, Phillips RJ, Shackel KA, McElrone AJ and Matthews MA (2011) Automated analysis of three-dimensional xylem networks using high-resolution computed tomography. New Phytologist 191(4), 1168–1179.

Burggraaf P (1972) Some observations on the course of the vessels in the wood of Fraxinus excelsior L. Acta Botanica Neerlandica 21(1), 32–47.

Cai J, Li S, Zhang H, Zhang S and Tyree MT (2014) Recalcitrant vulnerability curves: methods of analysis and the concept of fibre bridges for enhanced cavitation resistance. Plant, Cell & Environment 37(1), 35–44.

Carlquist S (1977) Ecological factors in wood evolution: a floristic approach. American Journal of Botany 64(7), 887–896.

Carmesin CF, Port F, Böhringer S, Gottschalk KE, Rasche V and Jansen S (2023) Ageing-induced shrinkage of intervessel pit membranes in xylem of Clematis vitalba modifies its mechanical properties as revealed by atomic force microscopy. Frontiers in Plant Science 14, 1002711.

Choat B, Badel E, Burlett R, Delzon S, Cochard H and Jansen S (2016) Noninvasive measurement of vulnerability to drought-induced embolism by X-ray microtomography. Plant Physiology 170(1), 273–282.

Choat B, Brodersen CR and McElrone AJ (2015) Synchrotron X-ray microtomography of xylem embolism in Sequoia sempervirens saplings during cycles of drought and recovery. New Phytologist 205(3), 1095–1105.

Choat B, Brodribb TJ, Brodersen CR, Duursma RA, López R and Medlyn BE (2018) Triggers of tree mortality under drought. Nature 558(7711), 531–539.

Choat B, Nolf M, Lopez R, Peters JM, Carins-Murphy MR, Creek D and Brodribb TJ (2019) Non-invasive imaging shows no evidence of embolism repair after drought in tree species of two genera. Tree Physiology 39(1), 113–121.

Dagan Z, Weinbaum S and Pfeffer R (1982) An infinite-series solution for the creeping motion through an orifice of finite length. Journal of Fluid Mechanics 115, 505–523.

Delzon S and Cochard H (2014) Recent advances in tree hydraulics highlight the ecological significance of the hydraulic safety margin. New Phytologist 203(2), 355–358.

Dixon HH and Joly J (1895) XII. On the ascent of sap. Philosophical Transactions of the Royal Society of London.(B.) (186), 563–576.

Duanmu JL and Chai WK (2025) Modelling innovation adoption spreading in complex networks. Applied Network Science 10(1), 10.

Dulamsuren C, Abilova SB, Bektayeva M, Eldarov M, Schuldt B, Leuschner C and Hauck M (2019) Hydraulic architecture and vulnerability to drought-induced embolism in southern boreal tree species of Inner Asia. Tree Physiology 39(3), 463–473.

Emory SF (1989) Principles of integrity-testing hydrophilic microporous membrane filters, part I. Pharmaceutical Technology 13(9), 68–77.

Esteban LG, Palacios P de, Gasson P, García-Iruela A, García-Fernández F and García-Esteban L (2024) Hardwoods: Anatomy and functionality of their elements—a short review. Forests 15(7), 1162.

Ewers FW, Ewers JM, Jacobsen AL and López-Portillo J (2007) Vessel redundancy: modeling safety in numbers. IAWA Journal 28(4), 373–388.

Gauthey A, Peters JM, Lòpez R, Carins-Murphy MR, Rodriguez-Dominguez CM, Tissue DT, Medlyn BE, Brodribb TJ and Choat B (2022) Mechanisms of xylem hydraulic recovery after drought in Eucalyptus saligna. Plant, Cell & Environment 45(4), 1216–1228.

González-Muñoz N, Sterck F, Torres-Ruiz JM, Petit G, Cochard H, Arx G von, Lintunen A, Caldeira MC, Capdeville G, Copini P et al (2018) Quantifying in situ phenotypic variability in the hydraulic properties of four tree species across their distribution range in Europe. PLoS One 13(5), e0196075.

Gostick J, Aghighi M, Hinebaugh J, Tranter T, Hoeh MA, Day H, Spellacy B, Sharqawy MH, Bazylak A, Burns A et al (2016) OpenPNM: a pore network modeling package. Computing in Science & Engineering 18(4), 60–74.

Guan X, Pereira L, McAdam SA, Cao KF and Jansen S (2021) No gas source, no problem: proximity to pre-existing embolism and segmentation affect embolism spreading in angiosperm xylem by gas diffusion. Plant, Cell & Environment 44(5), 1329–1345.

Held M, Jyske T and Lintunen A (2025) Coordinated_scaling_of_conduits_and_pits. Open data of publication “Conduit and pit dimensions scale in a coordinated way from the treetop to coarse roots in three boreal tree species”. GitLab, project number 4198.

Held M, Jyske T and Lintunen A (2026) Conduit and pit dimensions scale in a coordinated way from the treetop to coarse roots in three boreal tree species. Tree Physiology, tpag003.

Hethcote HW (2000) The mathematics of infectious diseases. SIAM Review 42(4), 599–653.

Hinrichsen H (2000) Non-equilibrium critical phenomena and phase transitions into absorbing states. Advances in Physics 49(7), 815–958.

Hölttä T, Cochard H, Nikinmaa E and Mencuccini M (2009) Capacitive effect of cavitation in xylem conduits: results from a dynamic model. Plant, Cell & Environment 32(1), 10–21.

Ingram S, Reischl B, Vesala T and Vehkamäki H (2024) Ruptures of mixed lipid monolayers under tension and supercooling: implications for nanobubbles in plants. Nanoscale Advances 6(15), 3775–3784.

Johnson S (2024) Epidemic modelling requires knowledge of the social network. Journal of Physics: Complexity 5(1), 01LT01.

Kaack L, Altaner CM, Carmesin CF, Diaz A, Holler M, Kranz C, Neusser G, Odstrcil M, Schenk HJ, Schmidt V et al (2019) Function and three-dimensional structure of intervessel pit membranes in angiosperms: a review. IAWA Journal 40(4), 673–702.

Kaack L, Weber M, Isasa E, Karimi Z, Li S, Pereira L, Trabi CL, Zhang Y, Schenk HJ, Schuldt B et al (2021) Pore constrictions in intervessel pit membranes provide a mechanistic explanation for xylem embolism resistance in angiosperms. New Phytologist 230(5), 1829–1843.

Karimi Z (2014) Systematic, Evolutionary and Functional Anatomy of Wood and Leaves in Betulaceae. PhD dissertation, Universität Ulm, Fakultät für Naturwissenschaften, Institut für Systematische Botanik und Ökologie.

Kenah E and Robins JM (2007) Second look at the spread of epidemics on networks. Physical Review E—Statistical, Nonlinear, and Soft Matter Physics 76(3), 036113.

Kivelä M, Arenas A, Barthelemy M, Gleeson JP, Moreno Y and Porter MA (2014) Multilayer networks. Journal of Complex Networks 2(3), 203–271.

Knipfer T, Brodersen CR, Zedan A, Kluepfel DA and McElrone AJ (2015) Patterns of drought-induced embolism formation and spread in living walnut saplings visualized using X-ray microtomography. Tree Physiology 35(7), 744–755.

Lintunen A and Kalliokoski T (2010) The effect of tree architecture on conduit diameter and frequency from small distal roots to branch tips in Betula pendula, Picea abies and Pinus sylvestris. Tree Physiology 30(11), 1433–1447.

Loepfe L, Martinez-Vilalta J, Piñol J and Mencuccini M (2007) The relevance of xylem network structure for plant hydraulic efficiency and safety. Journal of Theoretical Biology 247(4), 788–803.

Lu D, Smith-Martin CM, Muscarella R, Uriarte M and Zheng T (2025) A spatio-temporal model of embolism propagation in leaf vein networks. AoB PLANTS, plaf020.

Mantova M, Herbette S, Cochard H and Torres-Ruiz JM (2022) Hydraulic failure and tree mortality: from correlation to causation. Trends in Plant Science 27(4), 335–345.

McDowell N, Pockman WT, Allen CD, Breshears DD, Cobb N, Kolb T, Plaut J, Sperry JS, West A, Williams DG et al (2008) Mechanisms of plant survival and mortality during drought: why do some plants survive while others succumb to drought? New Phytologist 178(4), 719–739.

Mrad A, Domec JC, Huang CW, Lens F and Katul G (2018) A network model links wood anatomy to xylem tissue hydraulic behaviour and vulnerability to cavitation. Plant, Cell & Environment 41(12), 2718–2730.

Mrad A, Johnson DM, Love DM and Domec JC (2021) The roles of conduit redundancy and connectivity in xylem hydraulic functions. New Phytologist 231(3), 996–1007.

Nekovee M, Moreno Y, Bianconi G and Marsili M (2007) Theory of rumour spreading in complex social networks. Physica A: Statistical Mechanics and its Applications 374(1), 457–470.

Newman ME (2002) Spread of epidemic disease on networks. Physical Review E 66(1), 016128.

Newman ME (2003) The structure and function of complex networks. SIAM Review 45(2), 167–256.

Olson ME (2023) Imperforate tracheary element classification for studies of xylem structure-function relations. IAWA Journal 44(3-4), 439–464.

Pastor-Satorras R, Castellano C, Van Mieghem P and Vespignani A (2015) Epidemic processes in complex networks. Reviews of Modern Physics 87(3), 925–979.

Pereira L, Kaack L, Guan X, Silva LdM, Miranda MT, Pires GS, Ribeiro RV, Schenk HJ and Jansen S (2023) Angiosperms follow a convex trade-off to optimize hydraulic safety and efficiency. New Phytologist 240(5), 1788–1801.

Raponi S, Khalifa Z, Oligeri G and Di Pietro R (2022) Fake news propagation: a review of epidemic models, datasets, and insights. ACM Transactions on the Web (TWEB) 16(3), 1–34.

Rizi AK, Faqeeh A, Badie-Modiri A and Kivelä M (2022) Epidemic spreading and digital contact tracing: Effects of heterogeneous mixing and quarantine failures. Physical Review E 105(4), 044313.

Rosner S, Nöbauer S and Voggeneder K (2021) Ready for screening: fast assessable hydraulic and anatomical proxies for vulnerability to cavitation of young conifer sapwood. Forests 12(8), 1104.

Roth-Nebelsick A (2019) It’s contagious: calculation and analysis of xylem vulnerability to embolism by a mechanistic approach based on epidemic modeling. Trees 33(5), 1519–1533.

Sano Y, Morris H, Shimada H, Ronse De Craene LP and Jansen S (2011) Anatomical features associated with water transport in imperforate tracheary elements of vessel-bearing angiosperms. Annals of Botany 107(6), 953–964.

Schenk HJ, Steppe K and Jansen S (2015) Nanobubbles: a new paradigm for air-seeding in xylem. Trends in Plant Science 20(4), 199–205.

Sevanto S, Mcdowell NG, Dickman LT, Pangle R and Pockman WT (2014) How do trees die? A test of the hydraulic failure and carbon starvation hypotheses. Plant, Cell & Environment 37(1), 153–161.

Shah D and Zaman T (2011) Rumors in a network: Who’s the culprit? IEEE Transactions on Information Theory 57(8), 5163–5181.

Silva LdM, Bujnowski B, Pereira L, Miranda M, Schenk H and Jansen S (2025) Gas diffusion kinetics in relation to embolism formation and propagation in angiosperm xylem: a mini-review. Acta Horticulturae 1419, 123–134.

Silva LdM, Pereira L, Kaack L, Guan X, Pfaff J, Trabi CL and Jansen S (2024) The potential link between gas diffusion and embolism spread in angiosperm xylem: evidence from flow-centrifuge experiments and modelling. Plant, Cell & Environment 47(12), 4977–4991.

Silva LdM, Pfaff J, Pereira L, Miranda MT and Jansen S (2024) Embolism propagation does not rely on pressure only: time-based shifts in xylem vulnerability curves of angiosperms determine the accuracy of the flow-centrifuge method. Tree Physiology, tpae131.

Sperry JS and Hacke UG (2004) Analysis of circular bordered pit function I. Angiosperm vessels with homogenous pit membranes. American Journal of Botany 91(3), 369–385.

Stewart JJ, Allen BS, Polutchko SK, Ocheltree TW and Gleason SM (2025) Xylem embolism refilling revealed in stems of a weedy grass. Proceedings of the National Academy of Sciences 122(13), e2420618122.

Strati S, Patiño S, Slidders C, Cundall EP and Mencuccini M (2003) Development and recovery from winter embolism in silver birch: seasonal patterns and relationships with the phenological cycle in oceanic Scotland. Tree Physiology 23(10), 663–673.

Strickland C, Dangelmayr G, Shipman PD, Kumar S and Stohlgren TJ (2015) Network spread of invasive species and infectious diseases. Ecological Modelling 309, 1–9.

Tavares JV, Oliveira RS, Mencuccini M, Signori-Müller C, Pereira L, Diniz FC, Gilpin M, Marca Zevallos MJ, Salas Yupayccana CA, Acosta M et al (2023) Basin-wide variation in tree hydraulic safety margins predicts the carbon balance of Amazon forests. Nature 617(7959), 111–117.

Torres-Ruiz JM, Cochard H, Mencuccini M, Delzon S and Badel E (2016) Direct observation and modelling of embolism spread between xylem conduits: a case study in Scots pine. Plant, Cell & Environment 39(12), 2774–2785.

Tyree MT and Sperry JS (1989) Vulnerability of xylem to cavitation and embolism. Annual Review of Plant Physiology and Plant Molecular Biology 40(1), 19–36.

US EPA Office of Water (2005) Membrane filtration guidance manual. US Environmental Protection Agency.

Vergeynst LL, Dierick M, Bogaerts JA, Cnudde V and Steppe K (2015) Cavitation: a blessing in disguise? New method to establish vulnerability curves and assess hydraulic capacitance of woody tissues. Tree Physiology 35(4), 400–409.

Vespignani A (2012) Modelling dynamical processes in complex socio-technical systems. Nature Physics 8(1), 32–39.

Wason J, Bouda M, Lee EF, McElrone AJ, Phillips RJ, Shackel KA, Matthews MA and Brodersen C (2021) Xylem network connectivity and embolism spread in grapevine (Vitis vinifera L.) Plant Physiology 186(1), 373–387.

Watts DJ and Strogatz SH (1998) Collective dynamics of ‘small-world’ networks. Nature 393(6684), 440–442.

Wheeler JK, Sperry JS, Hacke UG and Hoang N (2005) Inter-vessel pitting and cavitation in woody Rosaceae and other vesselled plants: a basis for a safety versus efficiency trade-off in xylem transport. Plant, Cell & Environment 28(6), 800–812.

Yang D, Pereira L, Peng G, Ribeiro RV, Kaack L, Jansen S and Tyree MT (2023) A unit pipe pneumatic model to simulate gas kinetics during measurements of embolism in excised angiosperm xylem. Tree Physiology 43(1), 88–101.

Zhang Y, Pereira L, Kaack L, Liu J and Jansen S (2024) Gold perfusion experiments support the multi-layered, mesoporous nature of intervessel pit membranes in angiosperm xylem. New Phytologist 242(2), 493–506.

